# Epithelial-Mesenchymal Transition Induces FAM134A-mediated ER-phagy to Regulate Procollagen Secretion

**DOI:** 10.64898/2026.07.20.739610

**Authors:** Marisa Di Monaco, Ozlem Sen, Gabriella Saksono, Alessandro Vascon, Jie Ying Teo, Viktoria Brunner, Christian Schellhaas, Samuel Ulrich, Iolo Squires, Renata Filipe Soares, Marian Peteri, Jacob Corn, Jin Rui Liang

## Abstract

EMT is accompanied by extensive remodeling of the secretory pathway, yet how cancer cells maintain endoplasmic reticulum (ER) homeostasis during this transition remains unclear. Here we show that EMT activates ER-phagy as an adaptive mechanism to constrain secretory output. A genome-wide CRISPR activation screen unexpectedly identified EMT- and MET-associated transcription factors as opposing regulators of ER-phagy, and TGFβ-induced EMT robustly stimulated this pathway. Proteomic analysis revealed that ER-phagy regulates EMT-induced secretory protein degradation, including procollagens. Among FAM134 ER-phagy receptors, FAM134A uniquely mediated procollagen degradation through a non-canonical, LC3–independent mechanism. Loss of FAM134A diverted procollagen from degradation to secretion, resulting in a pro-invasive secretome, as conditioned media from FAM134A-deficient cells enhanced matrix invasion. These findings identify ER-phagy as a regulatory node that buffers EMT-associated secretory remodeling and reveal FAM134A as a context-specific suppressor of pro-invasive secretion.

## Introduction

The endoplasmic reticulum (ER) is one of the most abundant and multifunctional organelles, serving as the primary site for protein synthesis and folding, calcium storage, lipid biosynthesis and intracellular structural organisation ^1^. To ensure that these important roles are efficiently executed, the ER is dynamically regulated by three critical pathways - the ER associated degradation (ERAD), the unfolded protein response (UPR) and ER-specific autophagy (ER-phagy)- in response to changing intracellular and extracellular conditions ^2^. The specificity of ER-phagy is conferred by a set of ER-resident or ER-targeting autophagy receptors such as the FAM134 family proteins, CCPG1 ^4^, SEC62 ^3^, TEX264 ^4^ ^5^, RTN3 ^3^, ATL3 ^13^, CDK5RAP3 ^6,7^, and CALCOCO1 ^8^ which recruit the autophagy machinery to specific ER subdomains via their LC3-interaction region (LIR) or GABARAP-interaction motif (GIM).

While the mechanisms of ERAD and UPR have been extensively characterised, ER-phagy is comparatively underexplored and its physiological contribution towards cellular homeostasis remains poorly understood. ER-phagy mediates the selective turnover of ER subdomains through autophagy and is activated by stimuli such as nutrient deprivation, ER stress, or misfolded protein accumulation ^3–5,7–12^. Although classically viewed as a quality-control mechanism to clear defective proteins, accumulating evidence suggests that ER-phagy also remodels functional ER in response to changing cellular demands. Indeed, dysregulation of ER-phagy has been linked to various pathologies including neurodegeneration, metabolic disease, and cancer with poorly defined molecular mechanisms. Therefore, it is critical for us to understand how ER-phagy integrates quality control and adaptive ER remodeling to understand its physiological roles and pathological consequences.

In this study, we set out to perform a genome-wide CRISPRa gain-of-function screen which unexpectedly revealed a striking correlation between EMT status and ER-phagy activity. Epithelial-mesenchymal transition (EMT) is a fundamental biological process implicated in embryogenesis, wound healing and cancer metastasis ^13^. EMT is known to rewire the secretory pathway to boost production of various extracellular matrix proteins and signalling factors, thus placing substantial functional demands and stress on the ER. Although general autophagy is known to support cancer cell survival and metabolic adaptation, it remains unexplored how selective organelle-specific autophagy pathways, such as ER-phagy, are regulated during EMT or contribute to EMT-associated phenotypes.

We performed extensive characterisation of this novel EMT-induced ER-phagy pathway and demonstrated that EMT-driven synthesis of secretory proteins at the ER is coupled with enhanced ER-phagy. This process is mediated by the ER-phagy receptor, FAM134A, via a non-canonical, LC3-independent degradation pathway. Loss of FAM134A reroutes ER-phagy cargo toward extracellular secretion, thereby promoting cell invasion. These findings uncover a potential tumor-suppressive role for FAM134A and highlight a mechanistic link between EMT, ER-phagy, and metastatic potential.

## Results

### 1. Genome-wide CRISPR activation screen uncovers novel factors that regulate ER-phagy

We previously demonstrated that cDNA overexpression of most ER-phagy receptors robustly drives ER-phagy flux by increasing the recruitment of autophagic components to the ER surface ^6^. To explore additional factors that can similarly drive ER-phagy upon overexpression, we generated a clonal HCT116 colon cancer cell line that stably expresses a CRISPR-activation (CRISPRa) system consisting of a catalytically inactive Cas9 (dCas9) fused with the tripartite VP64-p65-Rta (VPR) transcriptional activators ^14^ (Fig 1A). We further introduced a modified version of a previously developed doxycycline-inducible ER-Autophagy Tandem Reporter (EATR) system (ssCALR-mCherry-eGFP-KDEL) into the HCT116-CRISPRa for fluorescence-activated cell sorting (FACS)-based quantitative measurement of ER-phagy (Fig 1B) ^5,15^. This reporter cell line is capable of driving transcriptional expression of several known ER-phagy receptors, including FAM134B ^9^, TEX264 ^4^ ^5^, and SEC62 ^3^ (Fig 1C, Supp Fig 1A). Importantly, CRISPRa-driven overexpression of these ER-phagy receptors significantly increases amino-acid starvation-induced ER-phagy as measured by FACS (Fig 1D).

**Figure 1.**
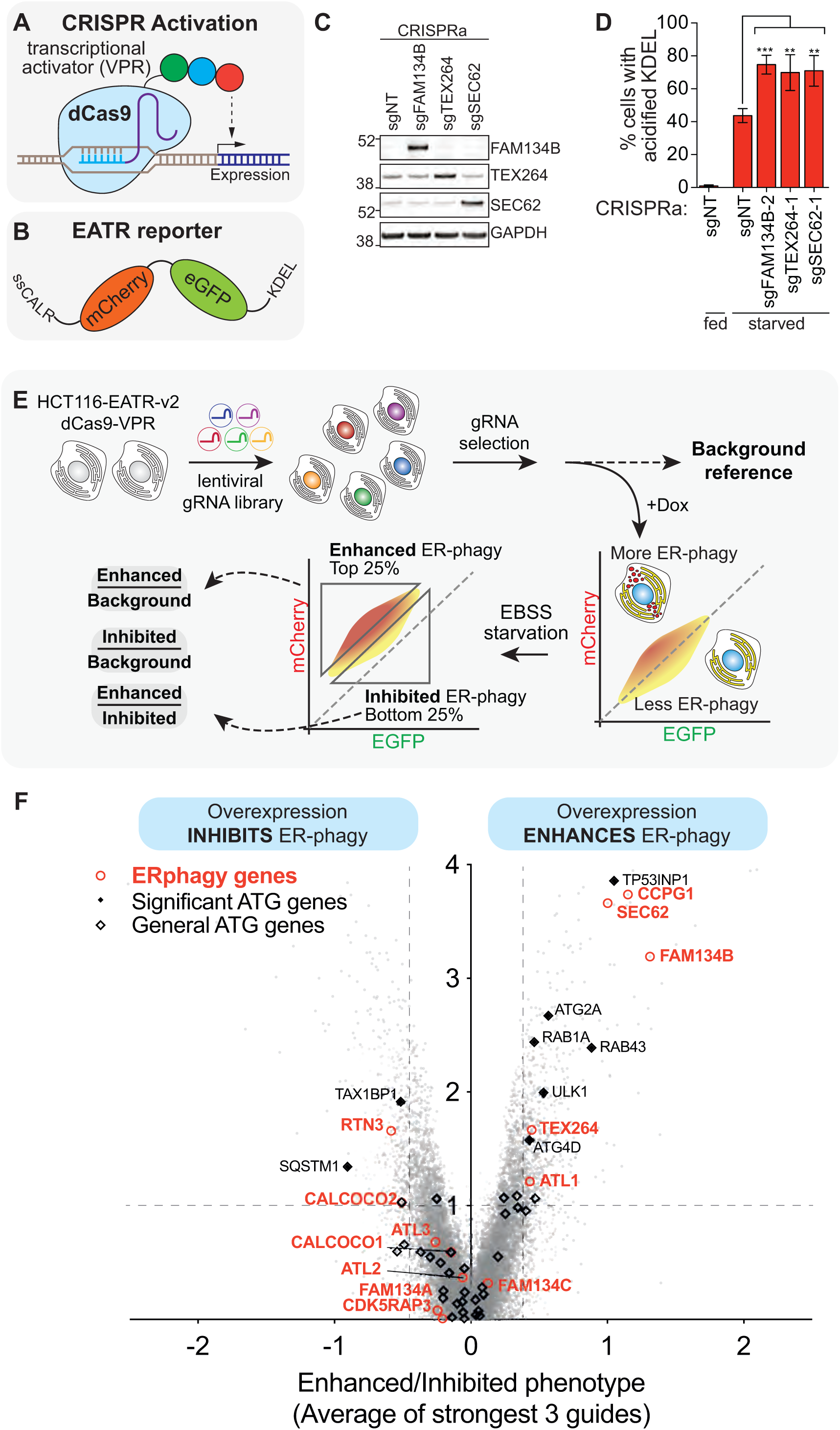
Genome-wide gain-of-function screen for ER-phagy factors. **(A)** A CRISPR activation (CRISPRa) system of dCas9-fused with VP64-p65-Rta is used to transcriptionally activate gene expression. **(B)** A tetracycline-inducible ER Autophagy Tandem Reporter (EATR) consists of an ER-targeting peptide from Calreticulin (ssCALR) fused with mCherry and GFP at the N-terminus, and ER retention signal (KDEL) fused to the C-terminus is used for FACS-measurement of ER-phagy flux. **(C)** HCT116-CRISPRa-EATR cells lentivirally transduced with the indicated sgRNAs (FAM134B, TEX264 and SEC62) were able to drive expression of the respective ER-phagy receptors. **(D)** CRISPRa-driven expression of the indicated ER-phagy receptors results in the expected ER-phagy enhancement quantified by EATR-FACS. **(E)** Workflow of the genome-wide EATR-FACS screening strategy to identify genes whose activation enhances or inhibits ER-phagy (see methods for detailed protocol). **(F)** CRISPRa screen data presented as volcano plot showing genes that upon overexpression either enhances (right side) or inhibits (left side) ER-phagy. Previously reported ER-phagy genes are annotated in red and known general autophagy genes are annotated in black.

**Supplementary Figure 1.**
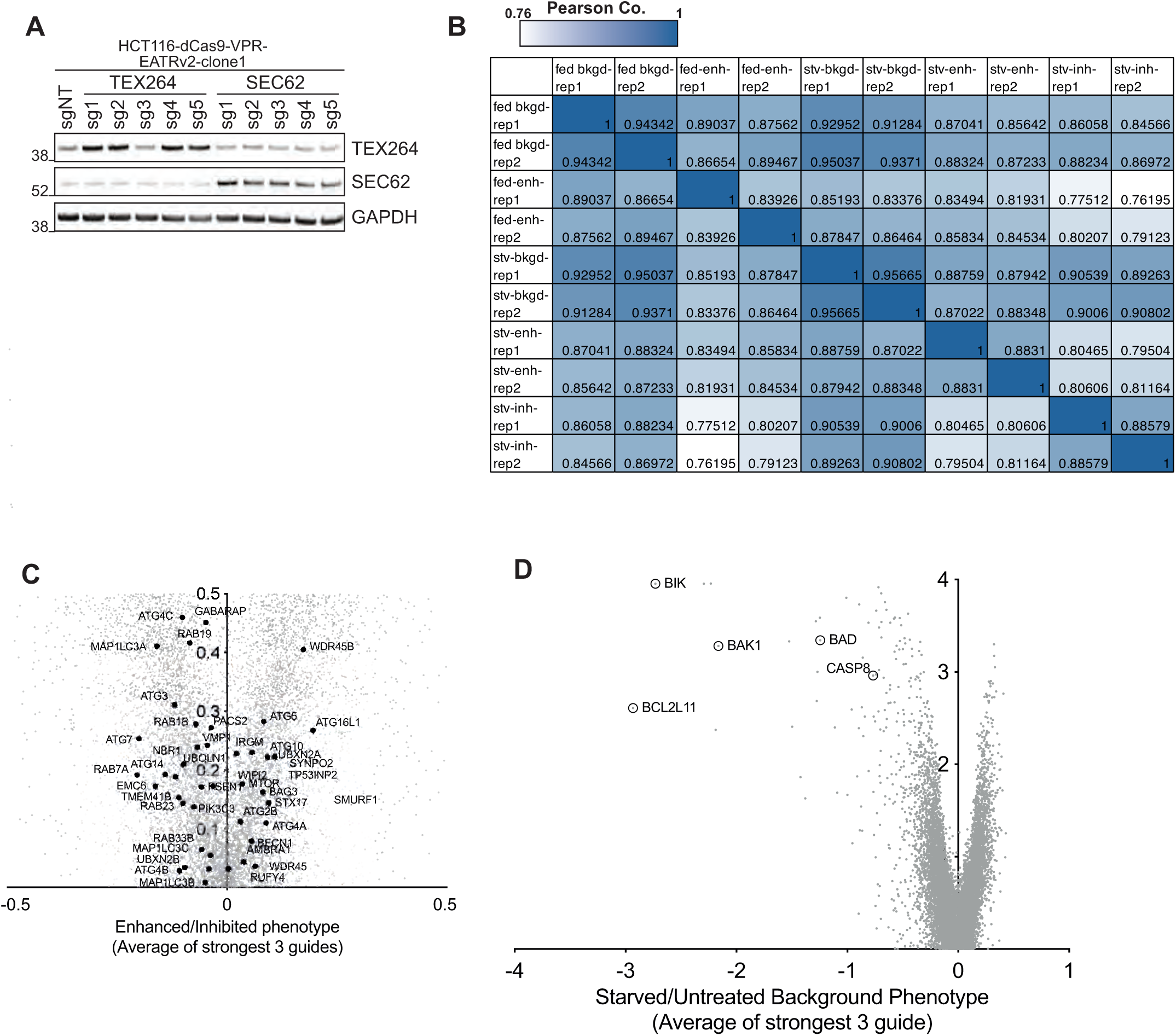
**(A)** A HCT116-CRISPRa clone was generated and validated to robustly activate gene expression using multiple guides targeting the promoter of TEX264 and SEC62. **(B)** Heatmap demonstrates biological reproducibility of the CRISPRa genome-wide screens. Two biological replicates were performed: represented as rep 1 & rep 2. (bkgd- unsorted background; enh- enhanced gate; inh - inhibited gate.) **(C)** Zoomed-in volcano plot of Fig 1F with general autophagy genes highlighted. **(D)** Volcano plot depicts genes that upon overexpression results in overall cell depletion between starved and untreated background samples. Known proapoptotic genes are annotated in red.

Next, we lentivirally transduced the cells with a genome-wide CRISPRa-V2 sgRNA library ^16^ (Fig 1E). We performed the screen either at basal (‘fed’) condition, or with amino-acid starvation (16hr incubation with Earl’s buffered saline solution; EBSS) to induce robust ER-phagy. Based on FACS measurement of mCherry and eGFP fluorescence intensities, we sorted for the top and bottom 25% of cells that correspond to gene overexpression that enhances and inhibits ER-phagy, respectively (Fig 1E).

We compared the unique sgRNA counts for each condition and observed strong correlation among their respective biological replicates, confirming the reproducibility of the screen (Supp Fig 1B). The lowest correlations were observed between the sorted populations corresponding to ‘‘enhanced’ and ‘inhibited’ ER-phagy, reflecting the divergence in genes that either promote or suppress ER-phagy. In line with previous reports, the screen robustly identified previously reported ER-phagy receptors including CCPG1, FAM134B and SEC62 as significant hits that increase ER-phagy upon overexpression (Fig 1F&G).

Notably, overexpression of most general autophagy-related genes did not increase ER-phagy, underscoring the specificity of the screen in capturing ER-phagy specific regulators (Fig 1F, Supp Fig 1C). We also note that a comparison of the starved versus basal (or ‘fed’) background samples revealed de-enrichment of multiple pro-apoptotic genes including BIK, BAK1, BAD, CASP8, and BCL2L11 upon starvation, exemplifying the robustness of the CRISPRa system in transcriptionally activating gene expression (Supp Fig 1D). Overall, these results demonstrate that our CRISPRa screen is technically robust and biologically specific, as evidenced by the successful recapitulation of known ER-phagy regulators, as well as identification of novel regulators of this pathway.

### 2. MAPK pathway activity and epithelial-to-mesenchymal transition status correlate with ER-phagy activity

Apart from previously reported ER-phagy receptors, Gene Ontology analysis of the top ‘activators’ and ‘inhibitors’ of ER-phagy revealed a surprising enrichment of genes involved in two pathways - the KRAS-MAPK pathway (Fig 2B-bold) and epithelial-mesenchymal transition (Fig 2A and B - black bars).

**Figure 2.**
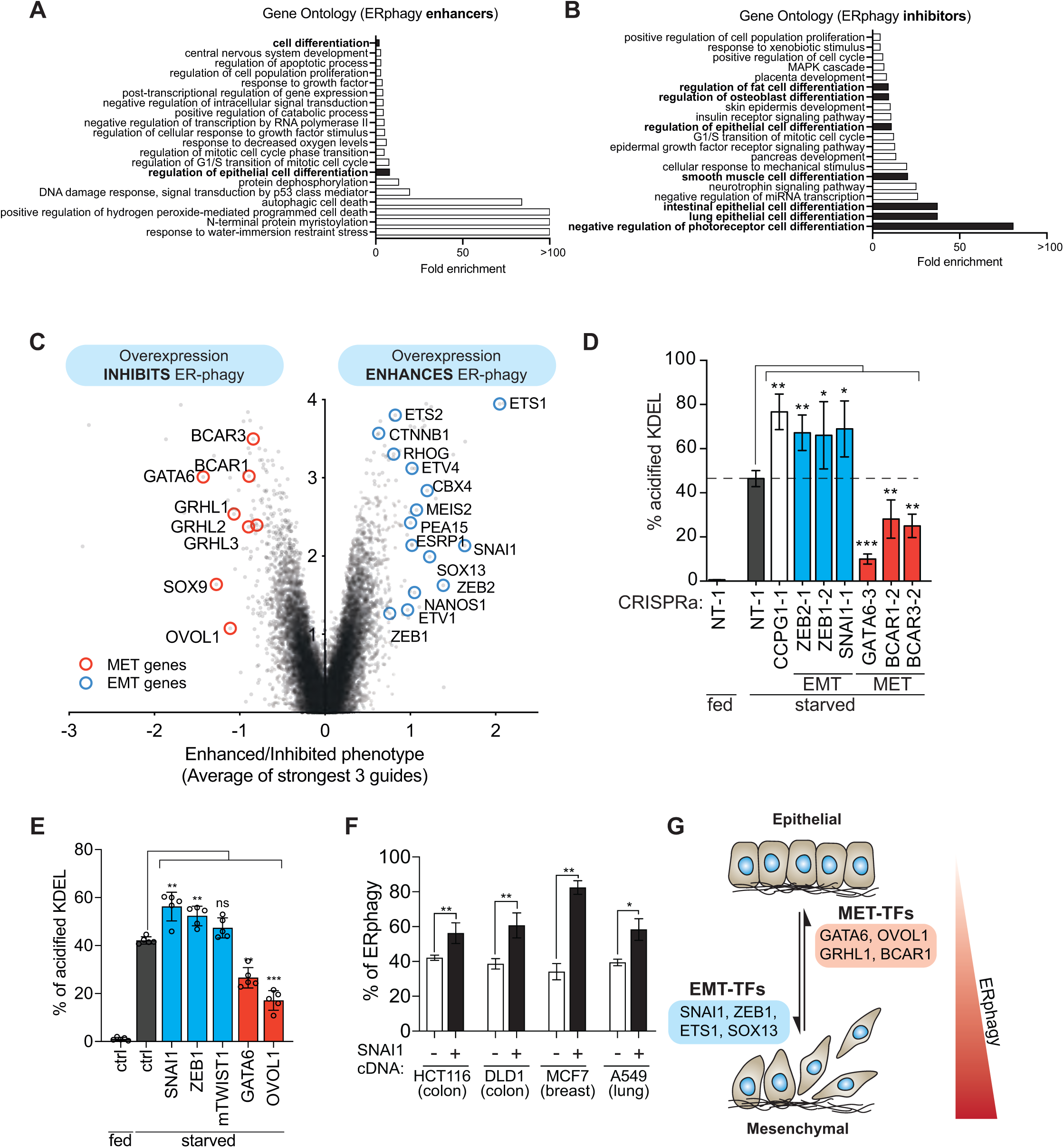
ERphagy positively correlates with EMT phenotype. **(A, B)** Gene ontology analysis on the top-enriched genes that activate (A) and repress (B) ER-phagy, respectively. Gene ontologies related to cell differentiation are bolded. **(C)** Volcano plot (same dataset as Fig 1F) is presented but with EMT and MET genes highlighted in blue and red, respectively. **(D)** Individual validation of screen results in HCT116-CRISPRa cells using sgRNAs confirmed that overexpression of EMT genes (in blue) enhances ER-phagy whereas overexpression of MET genes represses ER-phagy. Data represents ± SD of 3 biological replicates, paired t-test, ^∗^*p* ≤ 0.05, ^∗∗^*p* ≤ 0.01, ^∗∗∗^*p* ≤ 0.001. **(E)** Individual validation of screen results in HCT116-CRISPRa cells using cDNA overexpression confirmed that overexpression of EMT genes (in blue) enhances ER-phagy whereas overexpression of MET genes represses ER-phagy. Data represents ± SD of 4 biological replicates, paired t-test, ^∗^*p* ≤ 0.05, ^∗∗^*p* ≤ 0.01, ^∗∗∗^*p* ≤ 0.001 **(F)** Lentiviral overexpression of SNAI1 cDNA confirms that EMT enhances ER-phagy across various cell types measured using the EATR-FACS reporter. Data represents ± SD of 3 biological replicates for each cell line except DLD-1 (5 biological replicates), paired t-test, ^∗^*p* ≤ 0.05, ^∗∗^*p* ≤ 0.01, ^∗∗∗^*p* ≤ 0.001. **(G)** Our current working hypothesis suggests that the correlation between ER-phagy activity and EMT/MET status (regulated by the respective TFs) likely suggests that ER-phagy plays a contributing role during mesenchymal transition.

**Supplementary Figure 2.**
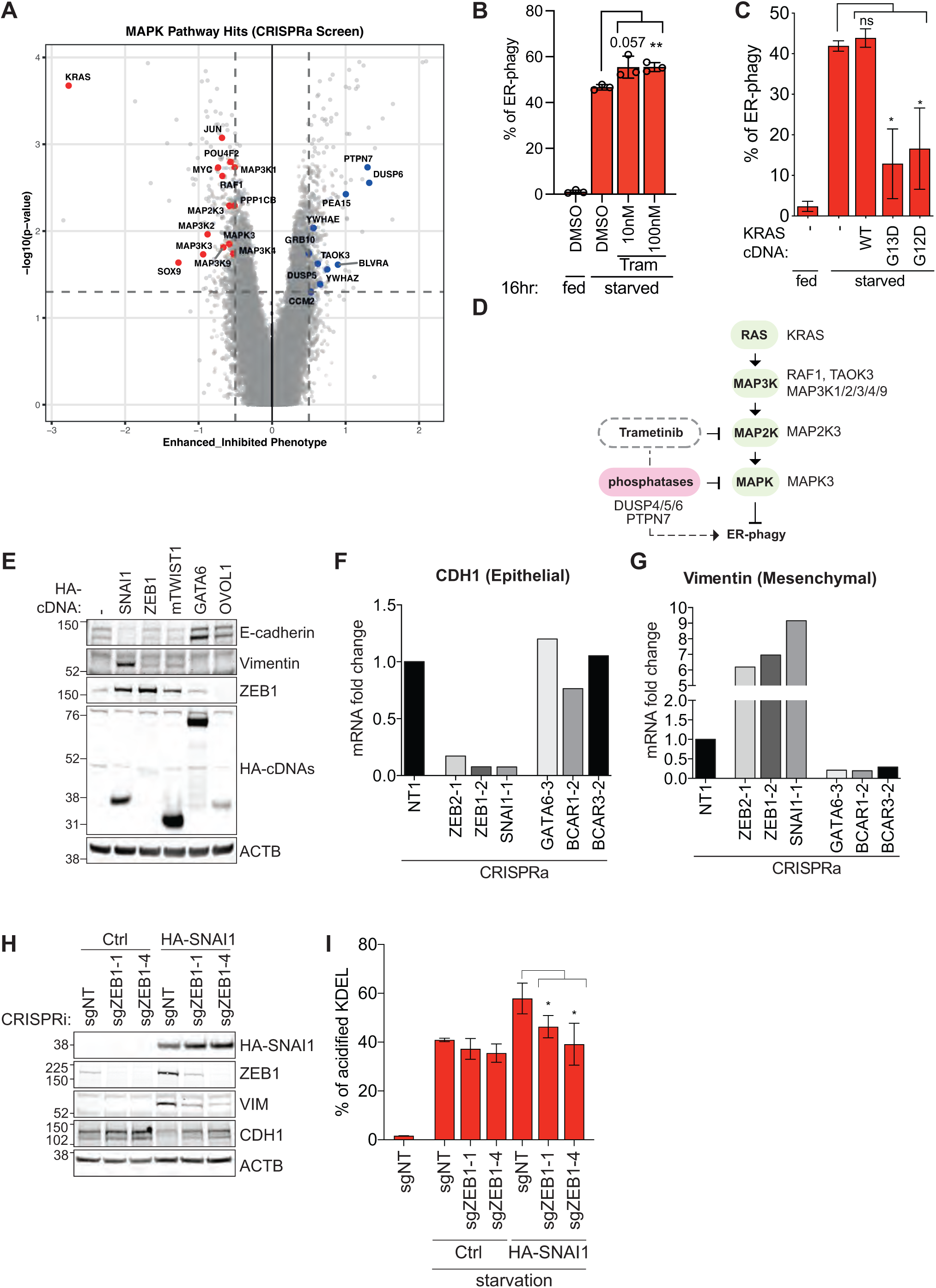
**(A)** Volcano plot (same dataset as 1F and 2C) is presented with genes of the KRAS-MAPK pathway highlighted, with those in red showing an inhibited phenotype and those in blue an enhanced phenotype. **(B)** Pharmacological inhibition of MEK1/2 with Trametinib shows enhanced ER-phagy after 16hr of EBSS treatment via FACS analysis. Data represents ± SD of 3 biological replicates, paired t-test, ^∗^*p* ≤ 0.05, ^∗∗^*p* ≤ 0.01, ^∗∗∗^*p* ≤ 0.001. **(C)** FACS analysis shows that overexpression of KRAS-G12D or G13D mutants, but not wild-type KRAS, robustly inhibits ER-phagy in HCT116 cells. Note that cells were only treated with EBSS for 6hr as overnight EBSS treatment led to excessive cell death in KRAS-G12D and G13D mutant cells. Data represents ± SD of 3 biological replicates, paired t-test, ^∗^*p* ≤ 0.05, ^∗∗^*p* ≤ 0.01, ^∗∗∗^*p* ≤ 0.001. **(D)** Schematic summarising the MAPK pathway-related genes identified from our CRISPRa screen. MAPK pathway activators led to repressed ER-phagy whereas MAPK pathway inhibitors (both genes and pharmacological inhibitors) enhance ER-phagy. **(E)** Western blot was used to validate the EMT status of cells used in Fig 2E. **(F, G)** Quantitative real-time PCR of cells used in Fig 2D confirmed that the indicated EMT and MET transcription factors indeed resulted in expected changes in CDH1 (epithelial marker) and Vimentin (mesenchymal marker). **(H, I)** CRISPRi-mediated knockdown of ZEB1 in SNAI1 overexpressing cells reversed the enhanced ER-phagy, shown both via Western blot **(H)** and EATR-FACS analysis **(I).** Data represents ± SD of 3 biological replicates, paired t-test, ^∗^*p* ≤ 0.05, ^∗∗^*p* ≤ 0.01, ^∗∗∗^*p* ≤ 0.001.

For the former, CRISPRa-mediated upregulation of MAPK pathway activators such as KRAS, RAF1, MAP3K1/2/3/4, MAPK3, and MAPK4, led to inhibition of ER-phagy (Supp Fig 2A). Conversely, overexpression of MAPK pathway repressors such as DUSP6 activates ER-phagy. Using pharmacological inhibition of MEK1/2 with Trametinib, we orthogonally validated that inhibition of the KRAS-MAPK pathway enhances ER-phagy (Supp Fig 2B) ^17^. Given that HCT116 cells harbor the KRAS-G13D mutation, we further showed that ER-phagy suppression requires oncogenic KRAS mutation, as only the overexpression of KRAS-G12D or G13D, but not wild-type KRAS robustly inhibited ER-phagy (Supp Fig 2C) ^17^. Overall, our data demonstrates that mutant KRAS-MAPK pathway activation inhibits ER-phagy (Supp Fig 2D). Since this observation is consistent with a recent report of similar observation whereby KRAS mutation disrupts ER-phagy in pancreatic ductal cells we opted to pursue another novel cluster of ER-phagy regulators ^18^.

The other striking cluster of regulators enriched in our genome-wide screen are transcription factors (TFs) that regulate epithelial-to-mesenchymal transition (EMT) and mesenchymal-to-epithelial transition (MET). EMT and MET are two fundamental and opposing pathways that dictate cell identity during development and are frequently co-opted during cancer metastasis (Fig 2C). This observation is both striking in terms of its specificity and directionality as canonical EMT-inducing transcription factors (EMT-TFs) such as SNAI1 ^19^, ZEB1/2 ^20^, and ETS1/2 ^21^ clustered exclusively as positive regulators of ER-phagy whereas well-established MET-inducing TF including OVOL1 ^22^, GRHL1/2/3 ^23^, and GATA6 ^24^ uniformly suppress ER-phagy. This suggests that the activation of ER-phagy might not merely be a byproduct of altered cell state, but could potentially be a regulated arm of the EMT program. Furthermore, the gain-of-function nature of our CRISPRa screen allowed us to identify redundantly acting gene paralogues, such as ETS1/2, SNAI1/2, ZEB1/2, BCAR1/3, and GRHL1/2/3 that may have overlapping functions during ER-phagy and would otherwise be masked in a CRISPRi or other loss-of-function approaches due to functional compensation.

We validated individual best-performing sgRNAs of several EMT- and MET-TFs from the CRISPRa screen data and confirmed that they indeed enhance and repress ER-phagy, respectively (Fig 2D). To rule out off-target effect of the sgRNAs, we also cloned the cDNA of several EMT- and MET-TFs, which further confirmed that exogenous expression of these genes phenocopied their CRISPRa phenotype (Fig 2E). Importantly, CRISPRa or cDNA overexpression of these EMT and MET-TFs resulted in the expected transition towards epithelial or mesenchymal status as assessed using CDH1 and Vimentin (mRNA and protein levels) as epithelial and mesenchymal markers, respectively (Supp Fig 2E-G).

Other studies have previously shown that SNAI1 transcriptionally upregulates ZEB1 ^25^, which we confirmed in HCT116 cells (Supp Fig 2H). Knockdown of ZEB1 in SNAI1 overexpressing cells reversed the enhanced ER-phagy, suggesting that SNAI1 acts upstream of ZEB1 to trigger ER-phagy (Supp Fig 2I). Using SNAI1 overexpression as a direct trigger for mesenchymal transition, we next asked if EMT-induced ER-phagy is a broadly conserved pathway. Due to EMT’s relevance in cancer metastasis, we assessed a panel of cancer cell types from different tissue sources, including colon cancer (HCT116 with KRAS G13D mutation and DLD1 with wild-type KRAS), breast cancer (MCF7 and MDA-MB-231), and lung cancer (A549). In all instances, SNAI1 overexpression results in enhanced ER-phagy (Fig 2F). Overall, our data suggest that activation of EMT- and MET-TFs antagonistically regulate ER-phagy (Fig 2G).

### 3. TGFβ-mediated mesenchymal transition also activates ER-phagy

To exclude possibility of artifacts due to artificial overexpression of EMT-TFs, we asked if physiological cytokines previously reported to trigger EMT such as TGFβ, IL6 and TNFα would also induce ER-phagy. At least in HCT116 and A549 cells, IL6 and TNFα do not induce mesenchymal transition or ER-phagy (Fig 3A). Consistent with the literature, we found that TGFβ robustly induces EMT in A549 cells as evidenced by the accumulation of several EMT-TFs such as SNAI1/2, ZEB1/2, TWIST1/2 (Fig 3A and Supp Fig 3A) and the decrease in CDH1 (Fig 3A). Concurrently, we observed the degradation of a major ER-phagy receptor, FAM134B and an increase in lysosomal cleavage of mCherry from the ER-phagy fluorescent reporter (CCER assay) ^15^, indicating that physiological activation of EMT induces ER-phagy (Fig 3A; Supp Fig 3A). Our model cell line, HCT116, has a loss-of-function mutation in TGFβ receptor II that renders it unresponsive to TGFβ treatment ^26^ (Fig 3A). Nevertheless, overexpression of SNAI1 overrides TGFβ-dependency and recapitulated the enhanced degradation of FAM134B and TEX264, indicating that ER-phagy is triggered by the TGFβ-SNAI1 pathway (Fig 3C) ^27^. Using EATR-FACS, we extended the TGFβ-induced ER-phagy observation to MCF7 cells (Fig 3C). We also observed TGFβ-driven loss of FAM134B in the non-transformed mouse mammary epithelial line NMuMG, demonstrating that EMT-induced ER-phagy may be a physiological and evolutionarily conserved response. (Supp Fig 3B).

**Figure 3.**
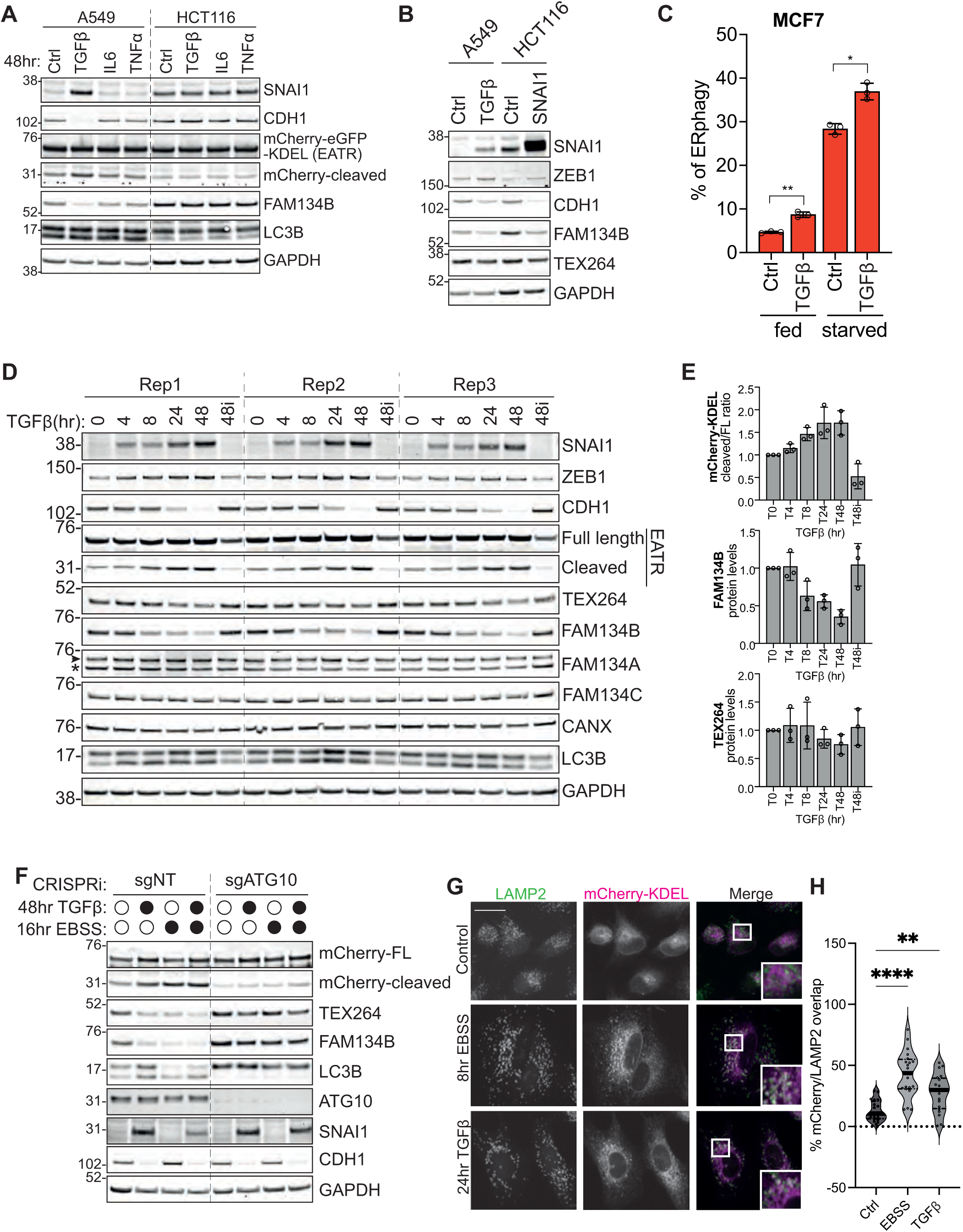
TGFβ stimulates EMT and induces ER-phagy. **(A)** HCT116 and A549 cells were treated with TGFβ (5ng/ml), IL6 (20ng/ml) and TNFa (10ng/ml) for 48 hr. Only TGFβ robustly induces EMT and ER-phagy in A549 lung carcinoma epithelial cells. HCT116 has a loss-of-function mutation in TGFβ receptor II, rendering it unresponsive to TGFβ treatment. **(B)** SNAI1 cDNA overexpression in HCT116 cells phenocopies downstream changes in TGFβ treatment in A549 cells, confirming that ER-phagy can be induced by the TGFβ-SNAI1 pathway. **(C)** EATR-FACS analysis shows an increase in ERphagy in response to TGFβ in MCF7 cells. Data represents ± SD of 3 biological replicates, paired t-test, ^∗^*p* ≤ 0.05, ^∗∗^*p* ≤ 0.01, ^∗∗∗^*p* ≤ 0.001. **(D, E)** Time-course assessment of EMT status and ER-phagy activity of A549-EATR cells upon TGFβ treatment. 48i indicates the use of 1µM SB505124 to inhibit the TGFβ signalling pathway, quantification of mCherry full length/cleaved, FAM134B and TEX264 shown in **(E). (F)** A549-CRISPRi cells with ATG10 knockdown were either treated with TFGβ, starved using EBSS, or co-treated with both. Blockage of the general autophagy pathway restores FAM134B and TEX264 levels. **(G)** Immunofluorescence of A549-ssCALR-mCherry-KDEL cells immunostained with LAMP2 to visualise the co-localisation of ER puncta within the lysosomes. Scale bar represents 20µm. Quantitation of the co-localisation of mCherry-KDEL puncta within the lysosomes after the indicated treatments. n = 20 cells, unpaired t-test, ∗∗p ≤ 0.01, ∗∗∗∗p ≤ 0.0001. **(H).**

**Supplementary Figure 3.**
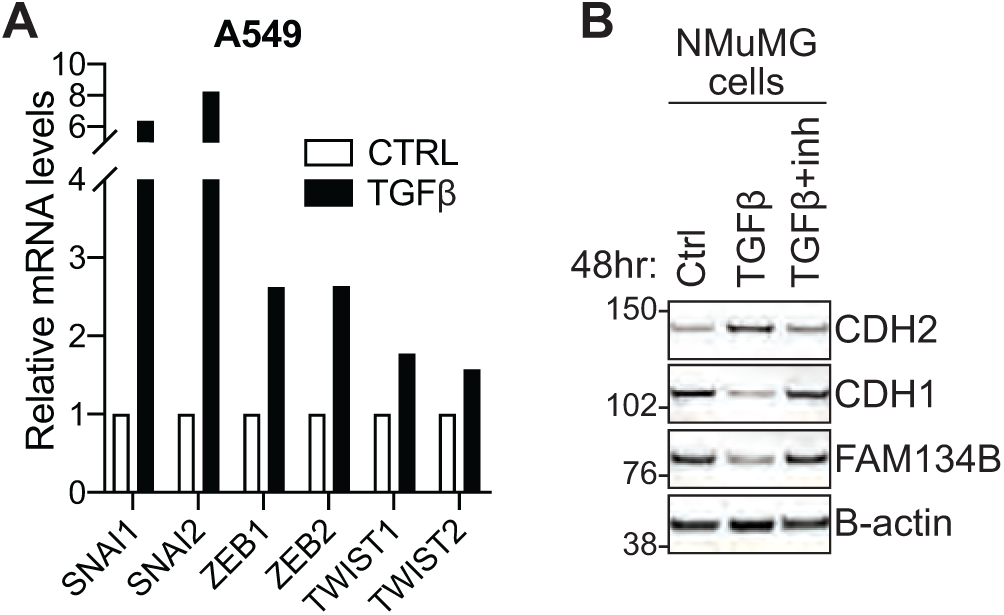
**(A)** Quantitative real-time PCR in A549 cells shows a robust upregulation of EMT genes upon treatment with TGFβ for 48 hr. Expression is relative to control conditions and normalised to β-Actin. **(B)** Western blot shows TGFβ-induced ER-phagy in a non-cancerous mouse mammary gland cell type, evidenced by the loss of FAM14B.

From hereon, we focused on using TGFβ treatment in A549 to physiologically induce EMT and we refer to the observed ER-phagy as EMT-induced ER-phagy. Over a 48-hour time course, we observed TGFβ treatment resulted in a progressive accumulation of the EMT transcription factors SNAI1 and ZEB1 (Fig 3D), accompanied by a loss of CDH1. As early as 8hr post-TGFβ treatment, ER-phagy was robustly activated, as evidenced by the loss of two ER-phagy receptors, FAM134B and TEX264, and to a lesser extent, the ER chaperone, CANX. We also detected the accumulation of lysosomally cleaved mCherry fragments from the ER-phagy reporter which increased over time. These changes were blocked with the use of a TGFβ-receptor inhibitor (SB505124; labelled as 48i), placing ER-phagy downstream of the TGFβ signalling pathway ^28^.

To confirm that the decrease of FAM134B and TEX264 is due to ER-phagy rather than EMT-associated transcriptional changes, we knocked down ATG10, a core component of the LC3-lipidation machinery. This led to the rescue of FAM134B and TEX264 protein levels upon TGFβ treatment (Fig 3F). We next used immunofluorescence microscopy to confirm ER engulfment within the lysosomes. TGFβ treatment resulted in a significant increase in ER puncta (ssCALR-mCherry-KDEL) within the lysosomes (LAMP2 staining) compared to untreated controls, indicating that a subset of ER is targeted for ER-phagy (Fig 3G). The extent of EMT-induced ER-phagy is slightly less than amino acid starvation-induced ER-phagy using EBSS (Fig 3H).

### 4. ER-phagy regulates abundance of EMT-induced secretory proteins

To understand the rationale of ER-phagy activation during EMT, we next sought to identify its cargoes. We performed total cell lysate mass spectrometry analysis comparing untreated versus TGFβ-treated A549 cells (Fig 4A). To capture proteins that further accumulate upon blockage of autophagy, we also included the lysosomal V-ATPase inhibitor, Concanamycin A (ConA). As expected, TGFβ treatment increased the abundance of many mesenchymal proteins (Fig 4B). We then filtered these upregulated proteins for those that were further elevated upon ConA treatment, as these are likely to be ER-phagy cargoes (Fig 4C and Supp Fig 4A). Among these, we observed several ER-synthesised secretory proteins such as SERPINE1/PAI-1, TIMP2, SPOCK1, ADAM19. Another prominent cluster consists of procollagens including COL4A1, COL4A2, COL5A1, and COL5A2 which showed marked accumulation upon ConA treatment (Supp Fig 4A). This was also accompanied by an increase in the ER-localised prolyl-4-hydroxylase, P4HA2, an enzyme involved in the proper folding of newly synthesised procollagen chains ^29^.

**Figure 4.**
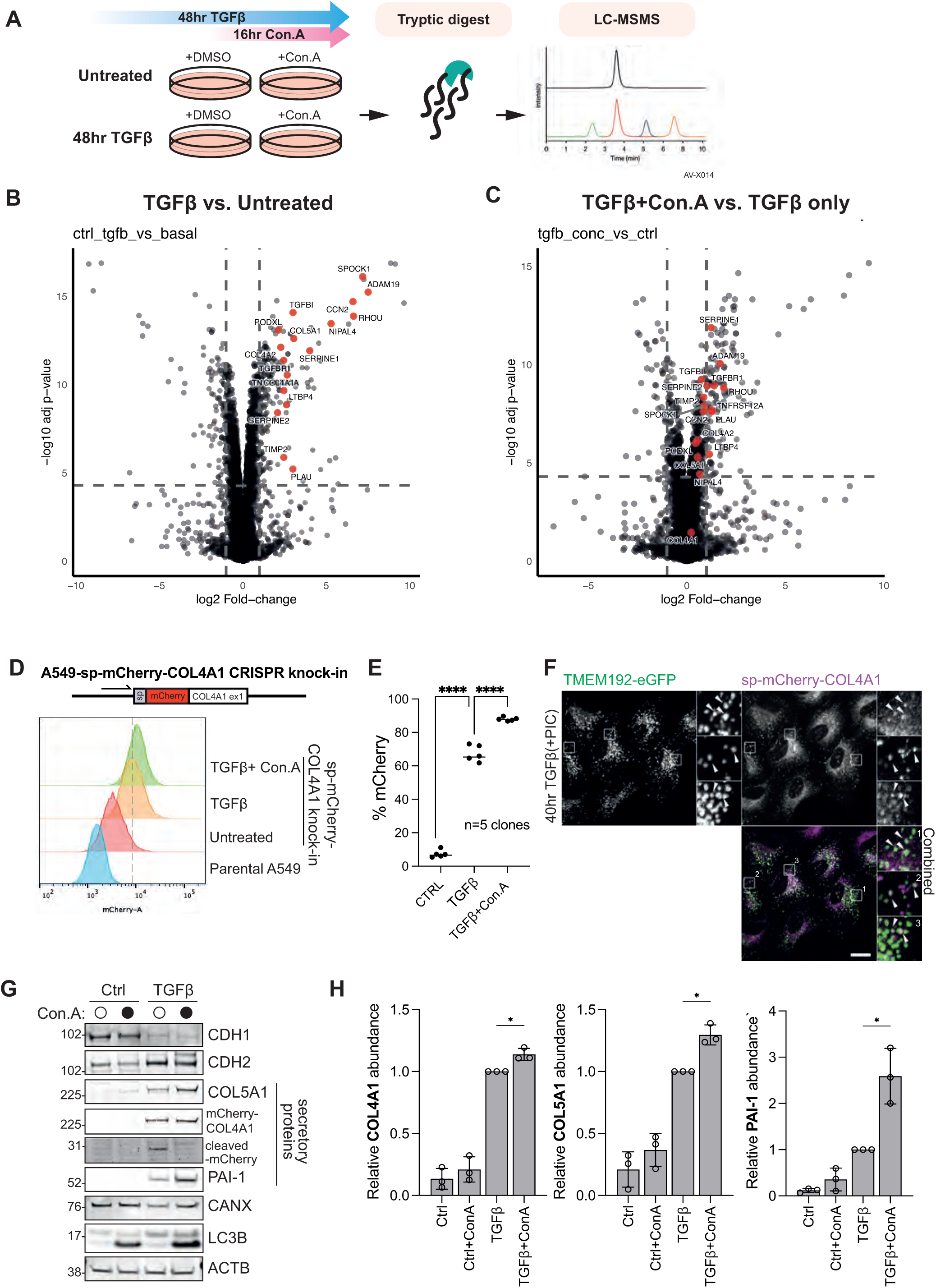
Pro-collagens are among the target substrates during EMT-induced ER-phagy. **(A)** Illustration of whole cell lysate mass spectrometry comparison of A549 cells treated with TGFβ for 48 hr. ConA treatment was introduced in the last 16hr to block lysosomal degradation of potential ER-phagy substrates. **(B,C)** Mass spectrometry data is presented as volcano plots first comparing untreated and TGFβ-treated A549 cells (B) and then TGFβ co-treatment with ConA versus DMSO (C). Annotated proteins indicate those that are upregulated upon TGFβ treatment and further accumulated upon ConA treatment. **(D)** A CRISPR/Cas9-knocked in mCherry-COL4A1 A549 cell line was generated to quantify procollagen autophagy. **(E)** 5 independent sp-mCherry-COL4A1 A549 clones using FACS showed consistent increase in signal upon 48 hr TGFβ treatment, which further increased upon Con.A co-treatment (last 24 hr). Data represents ±SD of 5 knock-in clones (as biological replicates), paired t-test, \*\*\*\**p* ≤ <0.0001. **(F)** Immunofluorescence images confirm that a fraction of mCherry-COL4A1 puncta colocalise with the lysosomal marker, TMEM192-eGFP upon treatment with 40hr TGFβ and PIC (last 16 hr). Scale bar represents 20µm. **(G,H)** Combined treatment of TGFβ (48 hr) and Con.A (last 24 hr) resulted in higher accumulation of COL4A1 (mCherry-tagged), COL5A1 and PAI-1 in A549 cells. Cleavage of mCherry by lysosomal proteases from the endogenously tagged mCherry-COL4A1 is also shown. Data represents ±SD of 3 biological replicates, paired t-test, ^∗^*p* ≤ 0.05.

**Supplementary Figure. 4.**
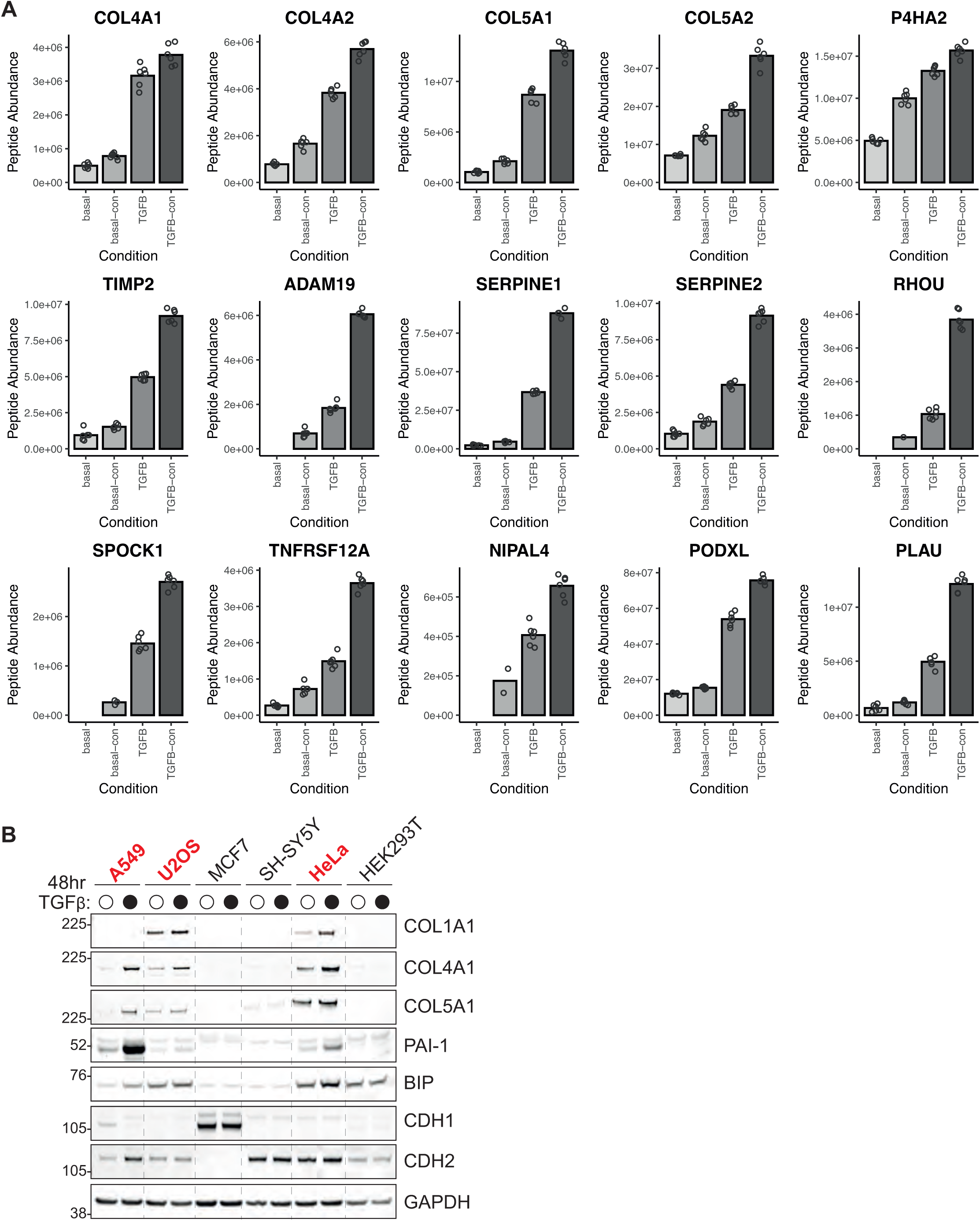
**(A)** Bar graphs show the raw peptide abundance of the potential ER-phagy cargoes across 6 biological replicates of whole cell lysate mass spectrometry analysis related to Fig 4A-C. **(B)** Comparison of several cancer cell lines showed an upregulation of COL1A1, COL4A1, COL5A1 and BIP in several TGFβ-responsive cell lines (highlighted in red; A549, U2OS and HeLa cells).

We validated the expression of some these secretory proteins across different cell types and found that procollagens and PAI-1 were robustly upregulated in TGFβ-responsive cell lines, including A549, HeLa and U2OS cells (Supp Fig 4B; annotated in red). In addition, we consistently observed increased levels of the ER chaperone BIP upon TGFβ treatment, suggesting that the increase in ER protein production contributes towards ER stress. We postulate that these secretory proteins, especially procollagens, might be targets of ER-phagy for several reasons. Firstly, all secretory proteins are synthesised and extensively modified in the ER ^30^. Second, up to 20% of nascent procollagens are known to be misfolded and are targets of ER-phagy via the FAM134 paralogues ^31,32^. Furthermore, disease mutations of procollagens that exacerbate their misfolding have been shown to trigger ER-phagy ^33,34^. It is therefore conceivable that ER-phagy might act as a quality control mechanism during mesenchymal transition to remove misfolded secretory proteins, or a regulatory system to control secretory output.

To visualise procollagen turnover via ER-phagy, we generated CRISPR/Cas9 knock-in A549 cells expressing endogenous mCherry-tagged COL4A1 which preserves its native promoter regulation and ER-targeting signal sequence (Fig 4D). This allows us to visualise endogenous COL4A1 localisation and measure its abundance in response to TGFβ treatment without artifact from aberrant overexpression. Using flow cytometry, we detected a robust increase in mCherry signal that is consistent with transcriptional upregulation of COL4A1. Importantly, co-treatment of cells with TGFβ and ConA resulted in higher mCherry signals, suggesting that a subset of mCherry-COL4A1 is normally targeted to the lysosomes (Fig 4D & E). We further characterised the subcellular localisation of mCherry-COL4A1 by immunofluorescence microscopy (Fig 4F). We observed that mCherry-COL4A1 localisation is consistent with both ER and puncta/vesicles. Importantly, some of the mCherry-COL4A1 puncta colocalised with the lysosomal marker TMEM192-eGFP, confirming that a fraction of these secretory proteins are delivered to the lysosome during EMT.

Using the mCherry-COL4A1 cells, we performed CCER assay and found that mCherry-COL4A1 was cleaved upon TGFβ and this was blocked upon ConA treatment (Fig 4G). In the same cell line, we also validated COL5A1 and PAI-1 accumulation upon TGFβ and ConA co-treatment by Western blotting (Fig 4G & H). Overall, our data support a model in which ER-phagy regulates secretory protein abundance during mesenchymal transition in response to the spike in ER protein synthesis demand.

### 5. FAM134A regulates pro-collagen turnover during EMT

The FAM134 family of ER-phagy receptors is known to mediate the turnover of misfolded procollagens at basal conditions ^34^. To determine if they are also responsible for procollagen degradation in response to TGFβ treatment, we knocked down FAM134A, B and C individually using siRNA. Interestingly, we found that only the loss of FAM134A resulted in a significant accumulation of COL4A1 and COL5A1 proteins (Fig 5A-C). Using qRT-PCR, we ruled out transcriptional upregulation as the contribution towards the increase in procollagen protein abundance upon FAM134A depletion (Supp Fig 5A). Within the same panel, we further probed and found that TGFβ treatment upregulates UPR markers including IRE1α, spliced XBP1 and BIP, consistent with our aforementioned observation that TGFβ lead to BIP protein accumulation (Supp Fig 5A & Supp Fig 4B). However, knockdown of FAM134A did not further elevate UPR response.

**Figure 5.**
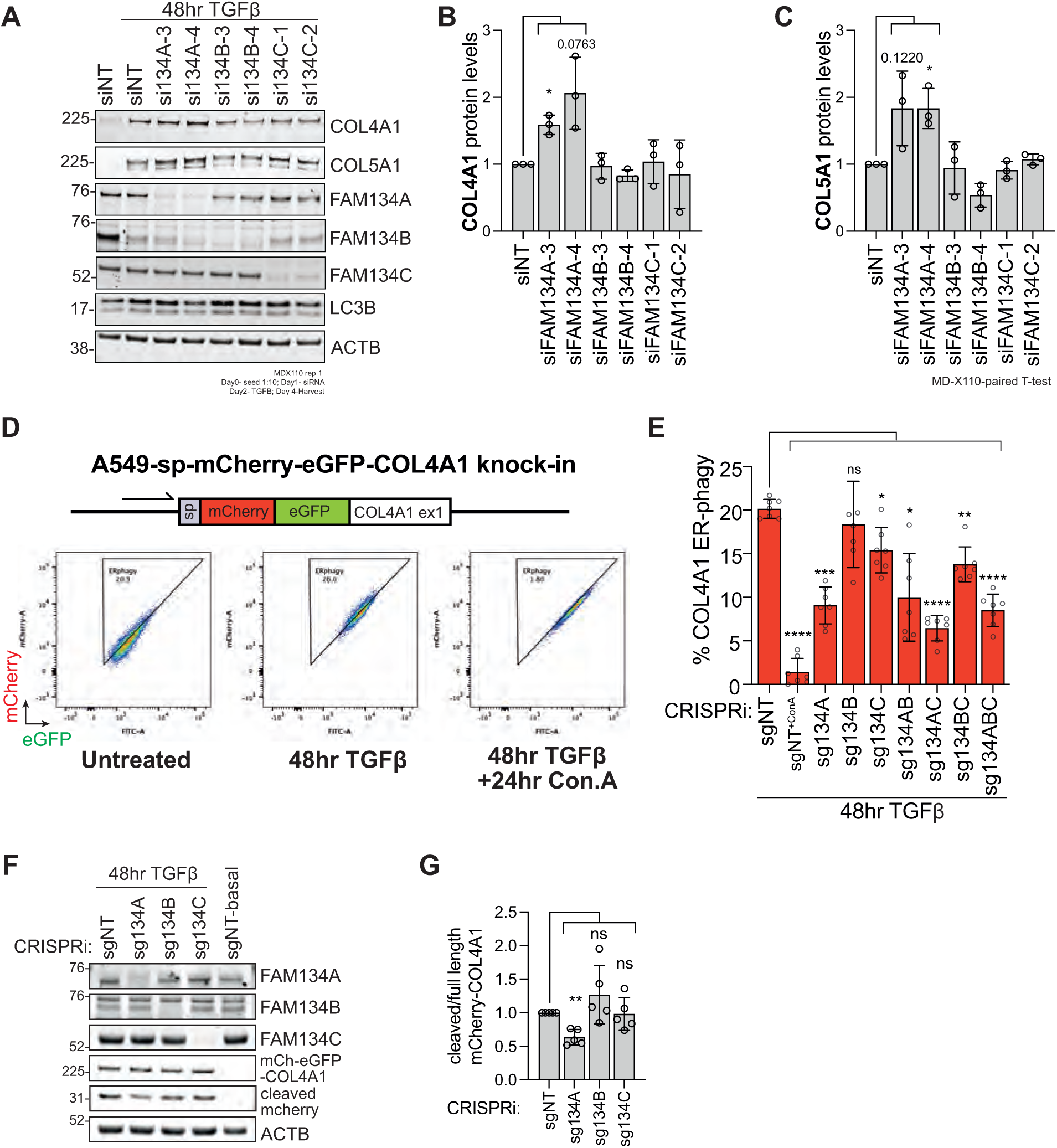
Knockdown of FAM134A blocks pro-collagen turnover. **(A)** Western blot data shows that only siRNA knockdown of FAM134A in A549 cells significantly increases the accumulation of COL4A1 **(B)** and COL5A1 **(C)**, relative to non-targeting siRNA control, siFAM134B and siFAM134C, upon 48 hr treatment with TGFβ. Data represents ±SD of 3 biological replicates, paired t-test, ^∗^*p* ≤ 0.05 **(D)** An endogenous mCherry-eGFP-tagged COL4A1 A549 cell line was generated using CRISPR/Cas9. EATR-FACS characterisation of this cell line showed a robust increase in mCherry and eGFP expression upon TGFβ treatment. Co-treatment with ConA regains eGFP signal, indicating that a fraction of COL4A1 is constitutively degraded via ER-phagy. **(E)** Combinatorial CRISPRi-based knockdown of FAM134A, B and C was performed in A549 cells. “% COL4A1 ER-phagy” indicates EATR-FACS measurement based on sp-Cherry-eGFP-COL4A1 (as opposed to the default ssCALR-mCherry-eGFP-KDEL) measured by FACS. Data represents ± SD of 7 biological replicates, one-way ANOVA and Dunnett’s post hoc test, ^∗^*p* ≤ 0.05, ^∗∗^*p* ≤ 0.01, ^∗∗∗^*p* ≤ 0.001, ∗∗∗∗p ≤ 0.0001, ns = *p* > 0.05. **(F, G)** Western blot data showing reduced mCherry-cleavage only with CRISPRi-mediated knockdown of FAM134A in A549-sp-mCherry-eGFP-COL4A1 cells upon TGFβ treatment. Data represents ±SD of 4 biological replicates, paired t test, ^∗∗^*p* ≤ 0.01, ns = *p* > 0.05.

**Supplementary Figure 5.**
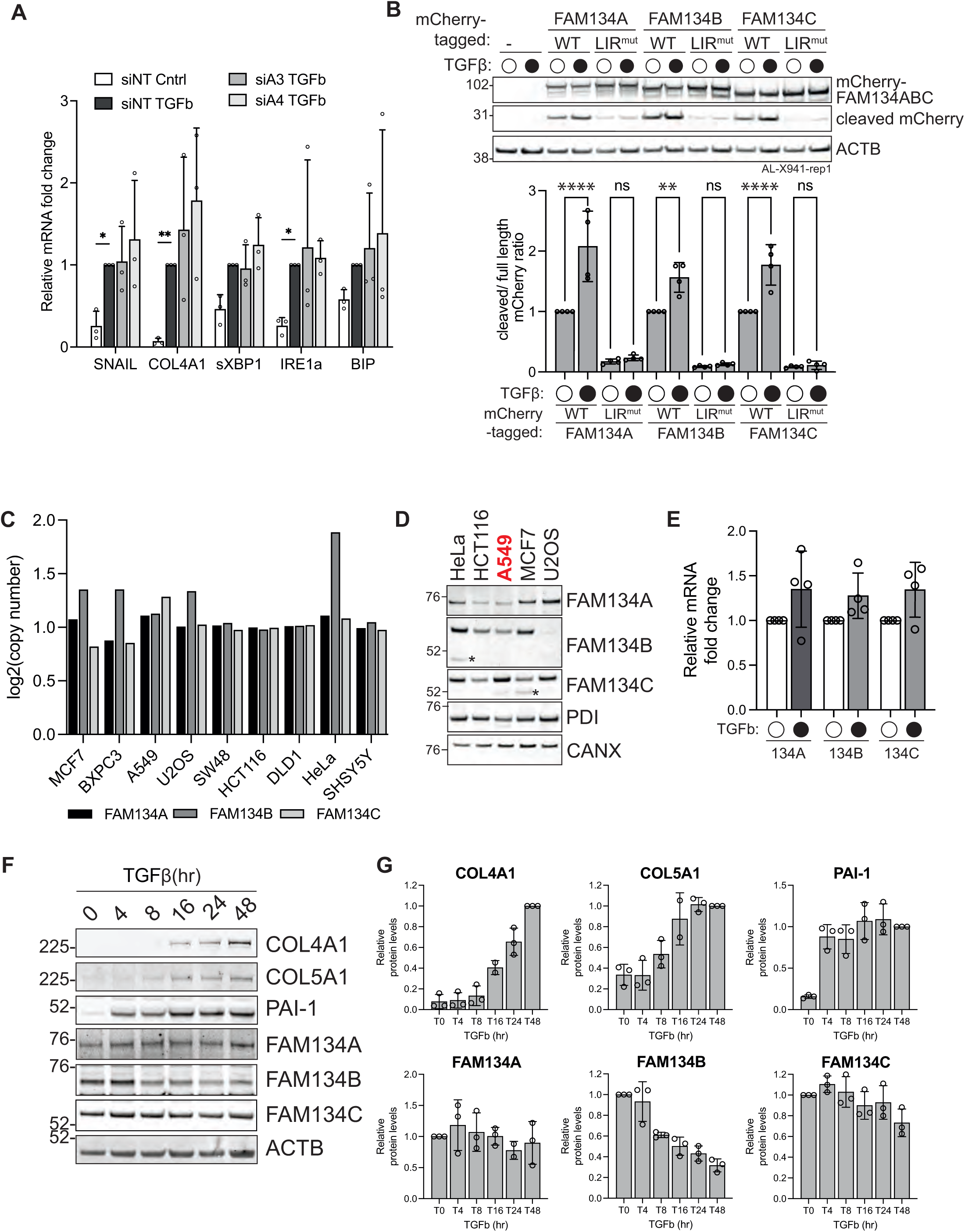
**(A)** Quantitative real-time PCR in A549 cells shows a robust upregulation of UPR genes sXBP1, IRE1a and BIP after 48 hr of TGFβ treatment, with no significant difference upon siRNA mediated knock down of FAM134A. Expression is relative to control conditions and normalised to GAPDH. Data represents ± SD of 3 biological replicates, two-way ANOVA and Dunnett’s post hoc test, ^∗^*p* ≤ 0.05, ^∗∗^*p* ≤ 0.01. **(B)** CCER analysis in A549 cells expressing mCherry-tagged FAM134A/B/C shows that all three paralogues undergo degradation during EMT-induced ER-phagy. Data represents ± SD of 4 biological replicates, Two-way ANOVA and Tukey’s multiple comparison post hoc test, ^∗∗^*p* ≤ 0.01, ∗∗∗p ≤ 0.001, ∗∗∗∗p ≤ 0.0001, ns = *p* > 0.05. **(C)** Bar graph depicts the absolute copy number of all three FAM134 proteins in commonly used cancer cell lines. Data obtained from public 24Q4 copy number database. **(D)** Relative protein levels of all three FAM134 paralogues across a panel of commonly used cancer cell lines, including A549 in red. **(E)** Quantitative real-time PCR in A549 cells shows that expression of FAM134A, FAM134B and FAM134C is not significantly changed in response to TGFβ treatment. **(F, G)** Time-course assessment of accumulation of COL4A1, COL5A1, FAM134A, FAM134B and FAM134C upon TGFβ treatment.

To attribute procollagen accumulation to ER-phagy inhibition, we again used CRISPR/Cas9 to knock-in a tandem mCherry-eGFP tag COL4A1 to monitor ER-phagy flux (Fig 5D). Using a similar gating strategy as previously established for EATR-FACS, we observed a strong increase in both mCherry and eGFP signals during TGFβ treatment, consistent with the transcriptional upregulation of COL4A1^12,15^. Importantly, co-treatment of these cells with TGFβ and ConA blocks eGFP loss while retaining mCherry signal compared to TGFβ treatment alone, indicating that a portion of the expressed COL4A1 is constitutively degraded via ER-phagy. This data is in line with the mCherry-COL4A1 CCER assay in Fig 4G.

We combined this mCherry-eGFP-COL4A1 reporter cell line with CRISPR interference (CRISPRi) to orthogonally knockdown FAM134A, B and C and explored their functions in COL4A1 turnover. Similar to our siRNA observation, only transcriptional repression of FAM134A resulted in robust inhibition of COL4A1 turnover (Fig 5E). To explore potential redundancy or synergy between the different FAM134 paralogues, we further performed combinatorial knockdowns and found that only conditions with FAM134A co-depletion resulted in strong inhibition of COL4A1 ER-phagy. We note that FAM134C depletion also resulted in a subtle but statistically significant inhibition (Fig 5E). Using the CCER assay, we orthogonally demonstrated that FAM134A knockdown resulted in a reduction in cleaved mCherry fragments, consistent with blocked ER-phagy (Fig 5F&G).

It was unexpected that FAM134A is the dominant ER-phagy reception under TGFβ stimulation, given that previous work reported higher ER-phagy flux driven by FAM134B and C compared to FAM134A ^34^. To determine whether all three FAM134 paralogues undergo degradation during EMT-induced ER-phagy, we expressed mCherry-tagged FAM134A/B/C (wild-type and LIR-mutant versions) and measure their turnover based on the accumulation of lysosomally-cleaved mCherry (Supp Fig 5B). As expected, the wild-type versions of all paralogues were constitutively degraded at basal conditions, consistent with their ability to trigger ‘auto’ ER-phagy as opposed to the corresponding LIR mutants. Upon TGFβ stimulation, turnover of all wild-type paralogues was further enhanced, tas evidenced by increased accumulation of cleaved mCherry products.

Transcriptomic analysis of the public 24Q4 dataset across a panel of cancer cell lines showed that all three FAM134 paralogues are expressed at comparable mRNA levels in A549 cells (Supp Fig 5C). When comparing the relative protein abundance of each FAM134 paralogue across commonly used cancer cell types, we observed that A549 cells exhibit the highest FAM134C, intermediate FAM134B, and lowest FAM134A expression (Supp. Fig 5D). Using qRT-PCR, we found that the transcript levels of all three FAM134 paralogues are not significantly altered upon TGFβ treatment, thus ruling out selective transcriptional upregulation of FAM134A as the reason for its specific requirement during EMT (Supp Fig 5E). Altogether, our data suggest that TGFβ-induced ER-phagy by FAM134A likely reflects its unique molecular function rather than its abundance levels when compared to the other FAM134 paralogues.

Lastly, we performed a time-course analysis following both collagen induction and FAM134B turnover. FAM134B was rapidly degraded as early as 8 hr after TGFβ treatment, whereas procollagen upregulation became pronounced after 16 hr (Supp Fig 4F&G). In contrast, FAM134A and FAM134C protein levels remained largely stable throughout. This observations could suggest a division of labour where FAM134B may mediate early TGFβ-induced ER-phagy of pre-existing substrates whereas FAM134A support a later stage ER-phagy of substrates such as procollagens that are transcriptionally induced by TGFβ.

### 6. FAM134A is able to mediate secretory protein turnover via a LIR-independent mechanism

To confirm the specificity of FAM134A’s role in COL4A1 ER-phagy, we reconstituted FAM134A cDNA in FAM134A knockdown cells. Re-expression of either wild-type or LC3-interacting region (LIR)-mutant FAM134A cDNA restored COL4A1 ER-phagy, consistent with a previous report of FAM134A’s non-canonical, LIR-independent, role in procollagen turnover (Fig 6A&B) ^34^.

**Figure 6.**
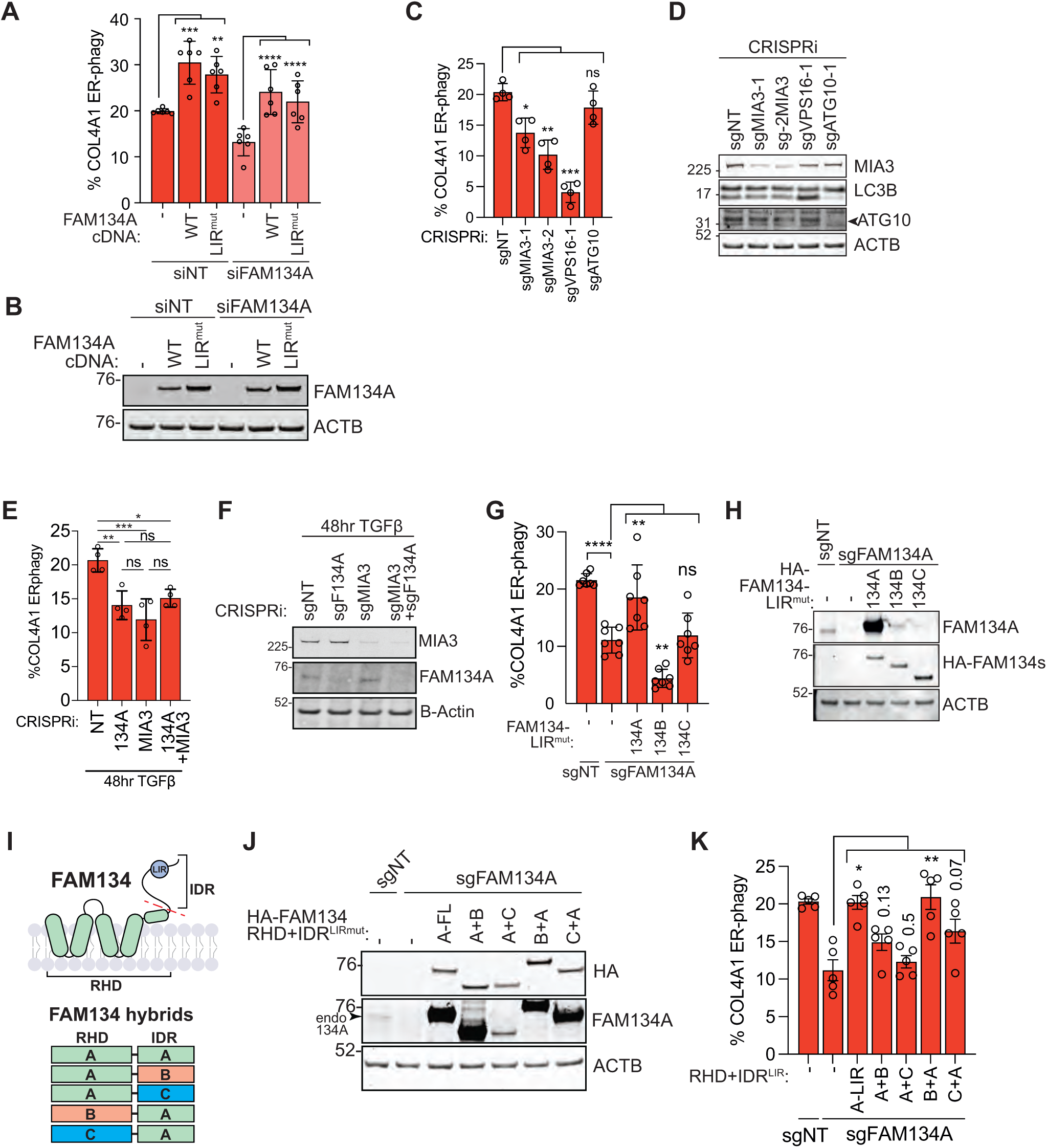
FAM134A regulates collagen turnover via a LIR-independent manner. **(A)** Reconstitution of wildtype and LIRmut cDNA re-expression in FAM134A knockdown cells rescyed COL4A1 ER-phagy based on EATR-FACS after 48 hr TGFβ treatment. Data represents ± SD of 3 biological replicates, two-way ANOVA and Dunnett’s multiple comparison test, ^∗∗^*p* ≤, ^∗∗∗^*p* ≤ 0.001, ∗∗∗∗p ≤ 0.0001. **(B)** Western blot validation of FAM134A knockdown and FAM134A cDNA re-expression using wild-type and LIR mutant (LIRMut) in A549 cells. **(C)** EATR-FACS analysis showing effects of the indicated knockdown on COL4A1 turnover. Data represents ± SD of 4 biological replicates, paired t-test *\*p* ≤ 0.05, ^∗∗^*p* ≤ 0.01, ^∗∗∗^*p* ≤ 0.001, ns = *p* > 0.05**. (D)** Western blot validation of CRISPRi-knockdown of MIA3, ATG10 and VPS16. **(E)** EATR-FACS analysis shows that double knockdown of MIA3 and FAM134A did not further inhibit COL4A1 degradation. Data represents ± SD of 4 biological replicates, paired t-test *\*p* ≤ 0.05, ^∗∗^*p* ≤, ^∗∗∗^*p* ≤ 0.001, ns = *p* > 0.05**. (F)** Western blot validation of MIA3 and FAM134A knockdown via CRISPRi. **(G)** Only the re-expression of FAM134A LIR^mut^ cDNA in FAM134A knockdown cells rescues COL4A1 ER-phagy as measured by EATR-FACS, Data represents ± SD of 6 biological replicates, paired t-test ^∗∗^*p* ≤ 0.01, ∗∗∗∗p ≤ 0.0001, ns = *p* > 0.05. **(H)** Western blot validation of expression of HA-FAM134 constructs**. (I)** A schematic illustrating the domain-swapping strategy for generating FAM134 hybrid constructs. **(J)** Western blot validation of the FAM134 hybrid cDNA re-expression in FAM134A knockdown cells. **(K)** FACS analysis shows that re-expression of hybrids containing FAM134A-IDR with FAM134B-RHD in FAM134A knock-down cells, can rescue COL4A1 turnover to similar levels as the FAM134A^LIRmut^. Data represents ± SD of 4 biological replicates, paired t-test *\*p* ≤ 0.05, ^∗∗^*p* ≤ 0.01, ns = *p* > 0.05.

**Supplementary Figure 6.**
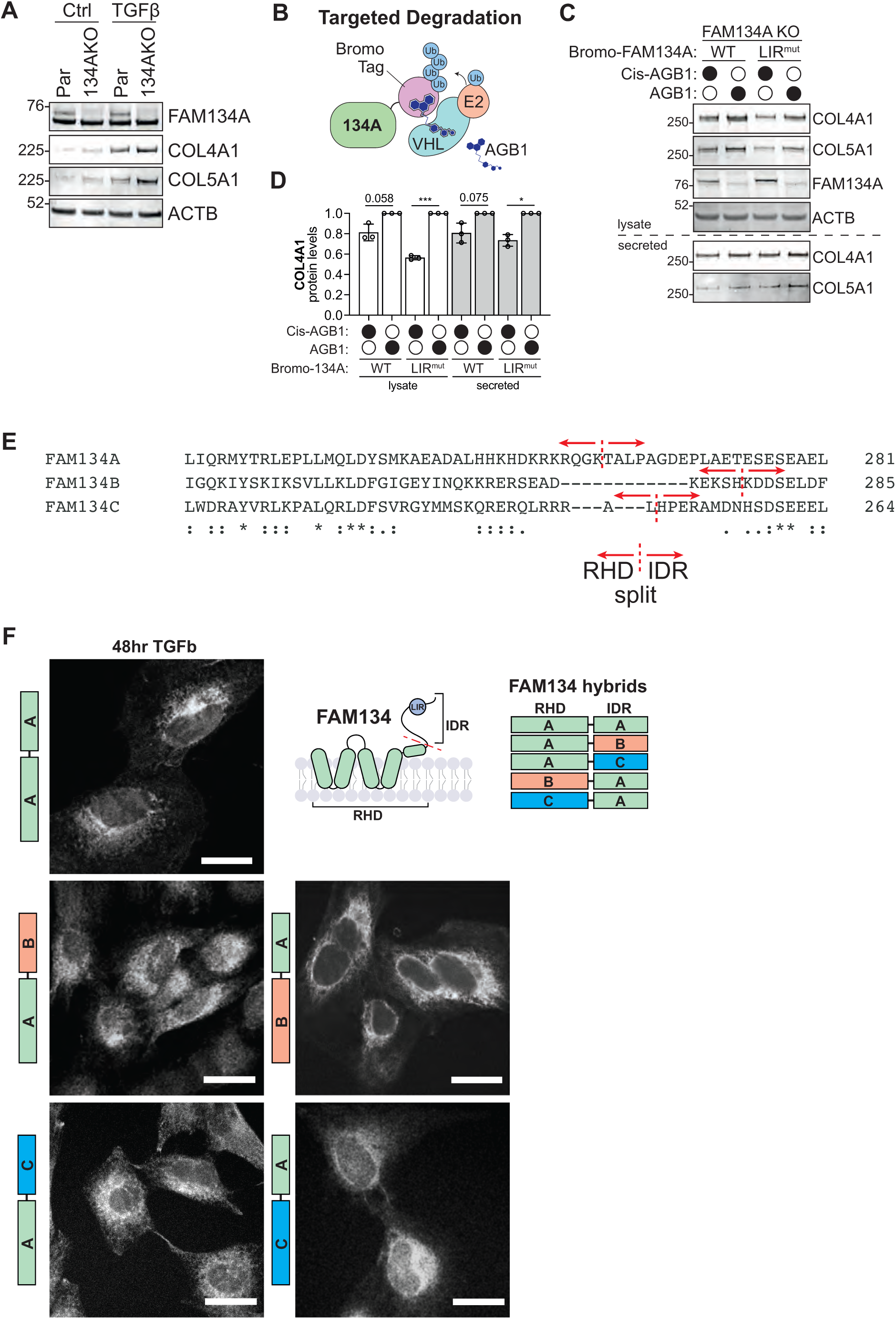
**(A)** Western blot data of FAM134A knockout (KO) in A549 cells shows accumulation of procollagens protein levels. **(B)** Schematic of Bromo-tagged FAM134A system, which allows inducible proteasomal degradation of FAM134A upon incubation with the small molecule AGB1. **(C,D)** Expression of Bromo-FAM134A wild-type and LIR mutant in FAM134A knock-out cells rescues both COL4A1 lysate and secreted protein levels. Data represents ± SD of 3 biological replicates, paired t-test, ^∗^*p* ≤ 0.05. **(E)** Protein sequence alignment of FAM134 paralogues at the region where the RHD and IDR were split. **(F)** Fluorescence microscopy was used (staining for HA-tagged FAM134 hybrids) to verify the correct ER-localisation of all FAM134 hybrid constructs. Scale bar represents 20µm.

We orthogonally validated the rescue experiments using a FAM134A knockout cell line expressing BromoTag-FAM134A cDNA (wild-type and LIRmut; Supp Fig 6A&B). This system allows acute, inducible proteasomal degradation of FAM134A upon incubation with the small molecule AGB1 ^35^. As a negative control, we used the inactive *cis*-AGB1 stereoisomer that does not engage VHL and therefore retains FAM134A. Consistent with the former rescue experiment (Fig 6A), inducible degradation of both BromoTag-FAM134A-WT and LIRmut using AGB1 resulted in accumulation of COL4A1 and COL5A1 (Supp Fig 6C&D).

We next explored the LIR-independent mechanism of COL4A1 degradation by perturbing distinct autophagy machineries. Knockdown of VPS16, a core component of HOPS/CORVET complex required for autophagosome-lysosome fusion, strongly inhibited COL4A1 turnover, confirming that procollagens are lysosomally degraded (Fig 6C&D)^36^. In contrast, knockdown of ATG10, the E2-like enzyme required for LC3 lipidation did not impair COL4A1 turnover despite robust inhibition of LC3 lipidation, indicating that LC3 conjugation is largely dispensable ^37^. This is in-line with our observation that FAM134A-LIRmut is able to restore procollagen turnover. Given that procollagens rely on the TANGO1/MIA3-COPII machinery for ER export^38^, and the COPII-associated non-canonical microautophagy has previously been reported to take place at ER exit sites ^31,39^, we asked whether this pathway also contributes to COL4A1 turnover during EMT. Indeed, MIA3 knockdown partially reduced COL4A1 degradation, suggesting that EMT-induced procollagen turnover engages, at least in part, a non-canonical autophagy mechanism associated with the TANGO1-COPII pathway and can take place independent of LC3 lipidation. Consistently, co-depletion of MIA3 and FAM134A did not further inhibit COL4A1 degradation compared with depletion of either gene alone, suggesting that the two proteins function in the same pathway regulating COL4A1 degradation (Fig 6E&F).

It is intriguing that FAM134B^LIRmut^ and FAM134C^LIRmut^ were unable to rescue ER-phagy despite sharing high structural homology with FAM134A (Fig 6G&H). Structurally, all three FAM134 paralogues share a conserved architecture consisting of an N-terminal reticulon-homology domain (RHD) and a C-terminal intrinsically disordered region (IDR), the latter being where the LIR resides (Fig 6K). We therefore sought to explore if FAM134A’s unique ability could be attributed to its RHD or the IDR domain by generating a series of domain-swapped FAM134 hybrid constructs fusing the RHD of FAM134B or C with the IDR of FAM134A, and vice versa (Fig 6I, Supp Fig 6E). In all cases, the LIR of each FAM134 paralogue was mutated to focus on exploring their LIR-independent roles. We verified by fluorescence microscopy that all FAM134 hybrids localised to the ER surface, thus ruling out mislocalisation as an explanation for the lack of ER-phagy rescue for any of the hybrids (Supp Fig 6F). When expressed in FAM134A knockdown cells, we found that re-expression of the FAM134B^RHD^-FAM134A^IDR-LIRmut^ and, to a weaker extent, FAM134C^RHD^-FAM134A^IDR-LIRmut^ were able to rescue COL4A1 turnover (Fig 6J&K). In contrast, none of the hybrids fused with FAM134A^RHD^ were able to rescue COL4A1 turnover. Overall, these observations suggest that the IDR of FAM134A likely interacts with unique factors other than the LC3/GABARAP family, or is subjected to specific post-translational modifications to mediate procollagen turnover.

### 7. FAM134A knockdown increases protein secretion and promotes cell invasion capacity

EMT drives cancer cell motility and invasion but it is unclear if EMT-induced ER-phagy contributes to these phenotypes. We conducted Kaplan-Meier survival analysis using the TCGA RNA-seq database accessed via GEPIA2 (Supp Fig 7A). Pan-cancer analysis revealed that higher expressions of FAM134A, B, and C were only modestly associated with improved overall survival. However, subtype-specific analyses highlighted a particularly strong correlation between higher FAM134A and FAM134B expression with improved survival outcome in kidney renal clear cell carcinoma (KIRC). These findings raise the possibility that FAM134A may exert a protective role in certain cancer types, potentially by modulating the secretory pathway to restrict cancer progression or metastasis.

**Fig. 7.**
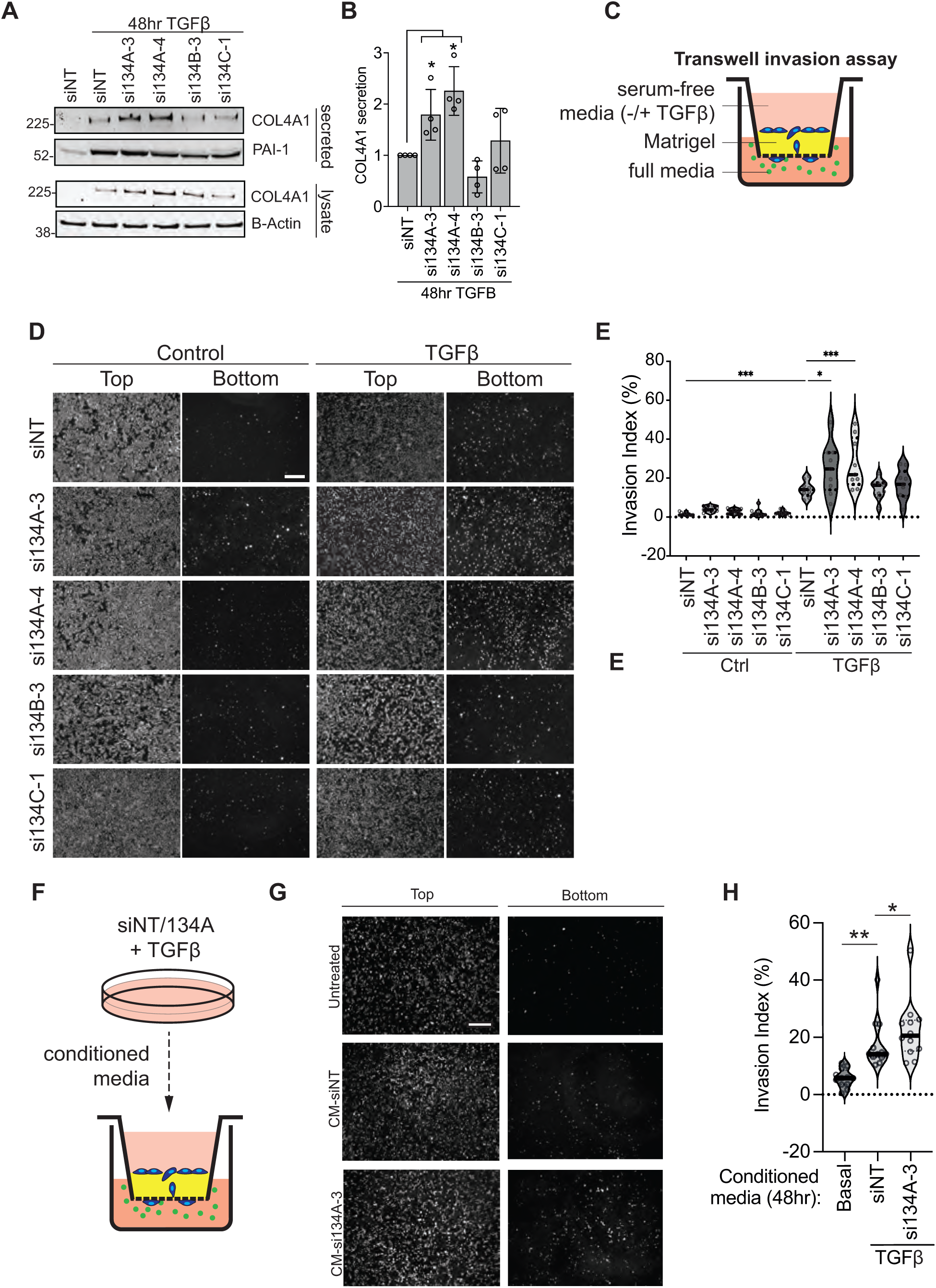
Increased secretion due to FAM134A knockdown promotes cell invasion capacity. **(A, B)** Conditioned media collected from A549 cells showed that knockdown of FAM134A significantly increases COL4A1 secreted protein levels following 48 hr TGFβ treatment. Data represents ±SD of 4 biological replicates, paired t-test, ^∗^*p* ≤ 0.05. **(C)** Invasion assay performed using Boyden chambers coated with Matrigel and cells treated with or without TGFβ for 48 hr. **(D)** Representative images of transwell assay for siRNA-mediated knockdown of FAM134A, FAM134B and FAM134C. Images represent cells at the top of the chamber (indicative of total cell number per chamber) and the bottom of the chamber (indicative of invasive cells). Scale bar represents 200µm. **(E)** Quantification of the invasion assay, with each data point representing an individual chamber across 3 biological replicates. siRNA-mediated knockdown of FAM134A results in a significant increase in invasion under TGFβ-treated conditions. One way ANOVA and Dunnett’s post hoc test, ^∗^*p* ≤ 0.05, ^∗∗∗^*p* ≤ 0.001**. (F)** Conditioned media collected from siFAM134A-A549 cells with and treated with TGFβ for 48 hr, was collected and plated on A549 parental cells inside Boyden chambers coated with Matrigel. **(G)** Following 48 hr incubation with TGFβ, the cells incubated with siRNA FAM134A knock-down conditioned media exhibited increased invasion. Scale bar represents 200µm. **(H)** Quantification of the invasion assay with each data point representing an individual chamber across 3 biological replicates. One way ANOVA and Dunnett’s post hoc test, ^∗^*p* ≤ 0.05, ^∗∗∗^*p* ≤ 0.001.

**Supplementary Figure 7.**
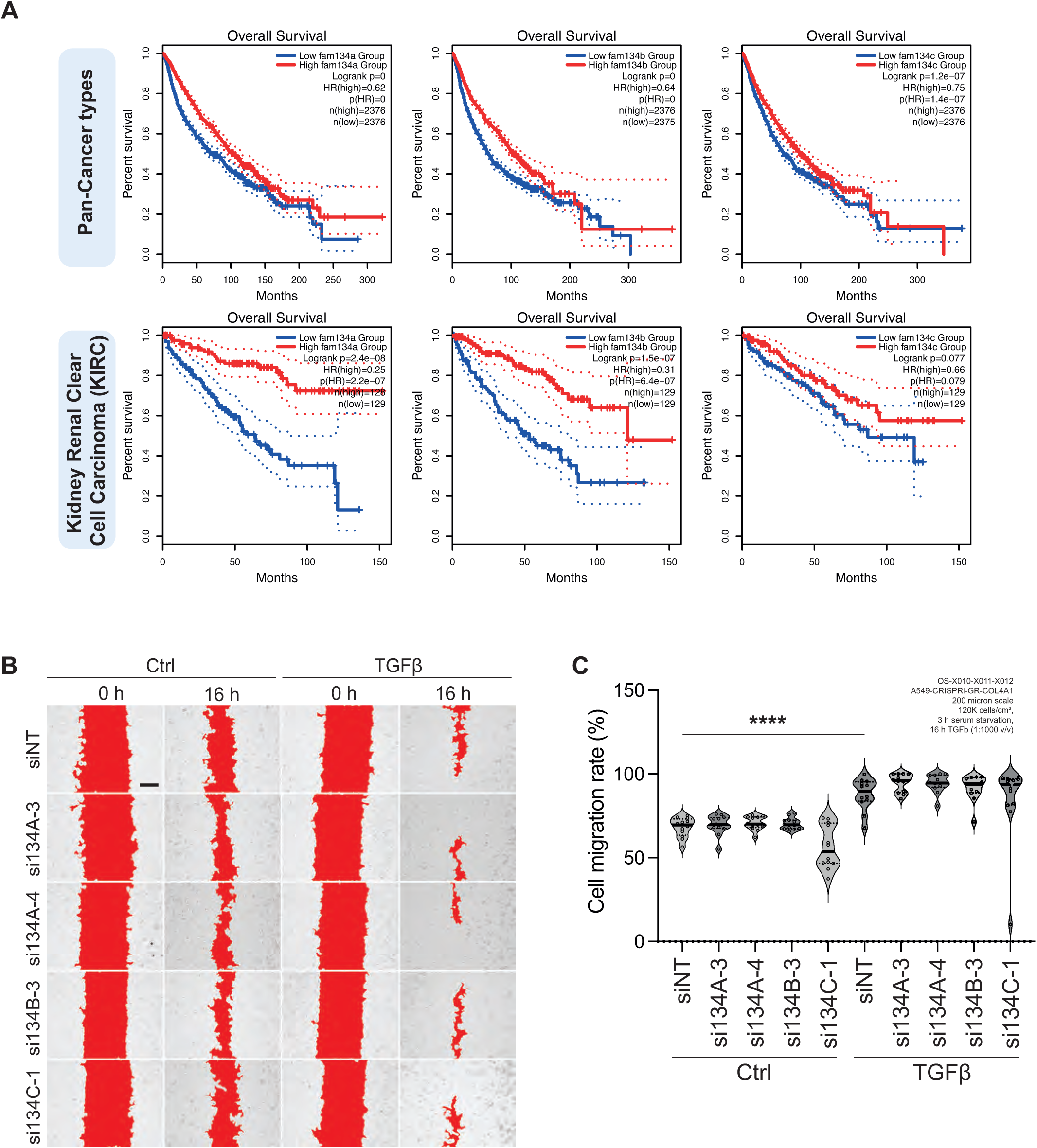
**(A)** Kaplan-Meyer survival curves illustrating the overall survival of patients stratified by FAM134A, B and C expressions based on RNA-seq data from The Cancer Genome Atlas (TCGA). Patients were grouped into high (top 25%, red), and low (bottom 25%, blue) expression cohorts using the quartile method. Survival analyses were performed using the GEPIA2 web server (http://gepia2.cancer-pku.cn). Results are shown for all cancer types (top row) and kidney renal clear cell carcinoma (KIRC, bottom row). Hazard ratios (HR), log-rank p-values, and sample sizes (n) for each group are indicated. Solid lines represent survival probabilities; dotted lines denote 95% confidence intervals. **(B)** Wound healing assay comparing cell migration after 16 hours TGFβ treatment showed that siRNA mediated knock-down of all three FAM134 paralogs does not impact cell migration. Representative images show wound closure at 0 hours and 16 hours. Scale bar represents 200µm. **(C)** Quantification of migration assay of four positions imaged per biological replicate (total 3 biological replicates). One-way ANOVA and Holm-Šidák post hoc test, ^∗∗∗∗^*p* ≤ 0.000.

To assess if these survival advantages might be attributed to ER-phagy, we explored the fate of the accumulated procollagen in the absence of FAM134A. To this end, we harvested conditioned media from A549 cells 48 hr post-TGFβ incubation and measured the amount of secreted COL4A1 upon each FAM134 protein knockdown (Fig 7A&B). We found that FAM134A depletion significantly increased the amount of secreted COL4A1 compared to FAM134B or C depletion, indicating that blocked ER-phagy reroutes COL4A1 from the lysosomes to the extracellular compartment. Inducible degradation of Bromo-FAM134A also recapitulated the increase in COL4A1 secretion (Supp Fig 6C&D; note that COL5A1 secretion is poorly detected by Western blotting and could not be reliably measured).

Increased secretion of collagens is known to drive ECM remodeling and invasion ^40^ ^41,42^.We first used a wound-healing assay to explore if FAM134A-mediated ER-phagy enhances cell migration in A549 cells. While TGFβ-induced mesenchymal transition increases overall wound closure, there was no significant difference between the non-targeting control and any of the FAM134 knockdown conditions (Supp Fig 7B). Next, we hypothesised that enhanced secretion as a result of ER-phagy blockage may promote invasion rather than migration. To explore this, we performed a Bodyen chamber invasion assay using matrigel-coated inserts to assess ECM degradation and invasion capability (Fig 7C). We found that TGFβ treatment significantly promoted cell invasion and this is further enhanced only in FAM134A knockdown conditions (Fig 7D-E).

We initially assumed that proteins destined for ER-phagy are likely misfolded and should thus be non-functional, even if secreted. To determine whether the enhanced invasion is due to increased secretion of functional proteins or strictly intracellular alterations due to FAM134A loss, we incubated wild-type A549 cells with conditioned media harvested from either control or FAM134A knockdown cells pre-treated with TGFβ for 48 hr (Fig 7F). Conditioned media from FAM134A knockdown cells significantly enhanced invasion of wild type A549 cells, suggesting that the effect is mediated by the secreted factors rather than intracellular changes (Fig 7G&H). Importantly, this suggests that the secreted factors are functionally capable of promoting cell invasion likely through remodeling the ECM.

## Discussion

Traditional loss-of-function screens have been instrumental in uncovering novel regulators of autophagy, yet they often suffer from high false-negative rates due to functional redundancies among gene paralogues ^6,43^. To overcome this limitation, we performed a gain-of-function CRISPRa screen in search of novel ER-phagy regulators. This approach not only identifies redundant paralogues within the same gene family but also enables interrogation of genes with low or no basal expression. The latter is particularly useful in overcoming cell/tissue type-specific gene expression barriers. Using this strategy, we uncovered the intrinsic oncogenic MAPK signaling pathway in HCT116 cells and the EMT/MET pathways as ER-phagy regulators.

EMT drives major transcriptional reprogramming that alters cell-cell adhesion, cell polarity and enhances migratory capacity. These changes are, in part, driven by a drastic increase in the synthesis and secretion of extracellular matrix (ECM) components and secretory factors such as matrix metalloproteases and procollagens. All secretory proteins are synthesis in the ER and this imposes substantial ER proteostatic stress. Our data demonstrate that this increase in secretory demand is coupled with elevated ER-phagy, likely serving as a homeostatic mechanism to regulate secretory output.

FAM134B and C have been implicated in myogenesis and neurodevelopment whereas the contribution of FAM134A remains elusive ^44,45^. A previous study showed that depletion of all FAM134 paralogues at basal state increases intracellular procollagen abundance without altering the total amount of procollagen secretion ^34^. During EMT, we observed that only FAM134A depletion led to significant increase in procollagen secretion. This discrepancy might reflect differences in experimental context. Prior studies primarily examined fibrillar collagen type I (COL1A1), whereas our work focuses on basement membrane–associated type IV and V collagens (COL4A1 and COL5A1), which might differ in their biosynthetic load, ER folding requirements, and supramolecular assembly challenges. Furthermore, prior studies explored basal secretion in cell types with constitutive procollagen expression (e.g. U2OS and MEFs), our studies were performed under TGFβ-stimulated conditions where secretory output and ER stress are significantly upregulated. It is conceivable that, under acute stimulated conditions, FAM134A-mediated ER-phagy becomes more critical to buffer excessive ER protein biosynthesis, and thus its function becomes more apparent than at basal state. Notably, depletion of FAM134B occurred as early as 8 hr post-TGFβ stimulation, preceding the induction and accumulation of COL4A1 and COL5A1, suggesting that differential temporal regulation might also contribute to the distinct requirement of each FAM134 paralogue.

Among the three paralogues, FAM134A harbours the largest cytoplasmic intrinsically disordered region (IDR) that might confer unique interactions with autophagy regulators distinct from FAM134B and FAM134C. Consistent with this, our domain-swapping analyses indicate that the IDR of FAM134A, rather than its reticulon-homology domain (RHD), is critical for its functional specificity. It is likely that FAM134A’s extended IDR may harbour distinct interaction motifs or post-translational modification sites that regulate its activity. Similar modifications have previously been described for FAM134B and FAM134C, which are regulated by CK2-mediated phosphorylation and AMFR-dependent ubiquitination ^46–48^.

A noncanonical procollagen autophagy pathway mediated by MIA3-COPII via the ER exit site has previously been reported, although the functional requirement of LC3 was not explored in that case^31,39^. Consistent with the ATG10- and LC3-independent role of FAM134A in COL4A1 turnover, we found that COL4A1 degradation requires, at least in part, the MIA3–COPII machinery. Therefore, FAM134A might facilitate multiple procollagen degradative pathways- both LC3-dependent and MIA3/COPII-dependent degradations - to buffer the surge in procollagen synthesis.

Our data further indicates that loss of FAM134A reroutes procollagen destined for degradation towards secretion, thereby increasing cancer cell invasion capability. Importantly, the COL4A1 initially targeted for ER-phagy are not misfolded beyond repair. Instead, they are likely to be partially-misfolded intermediates that are still biologically functional to promote invasion. Similar degradation of partially functional proteins has been observed in pancreatic ductal carcinoma where ER-retained MHC-I proteins targeted for lysosomal degradation remain functional when rerouted to the plasma membrane ^49^. Of note, although COL4A1 emerges as the primary substrate in our system, it remains unclear whether FAM134A also influences the secretion of other invasion-promoting secretory proteins. FAM134A-mediated ER-phagy may therefore serve a fine-tuning role to limit the burden of excessive secretion (Fig 8). This would position FAM134A as a brake on ECM remodeling and invasion.

**Fig. 8.**
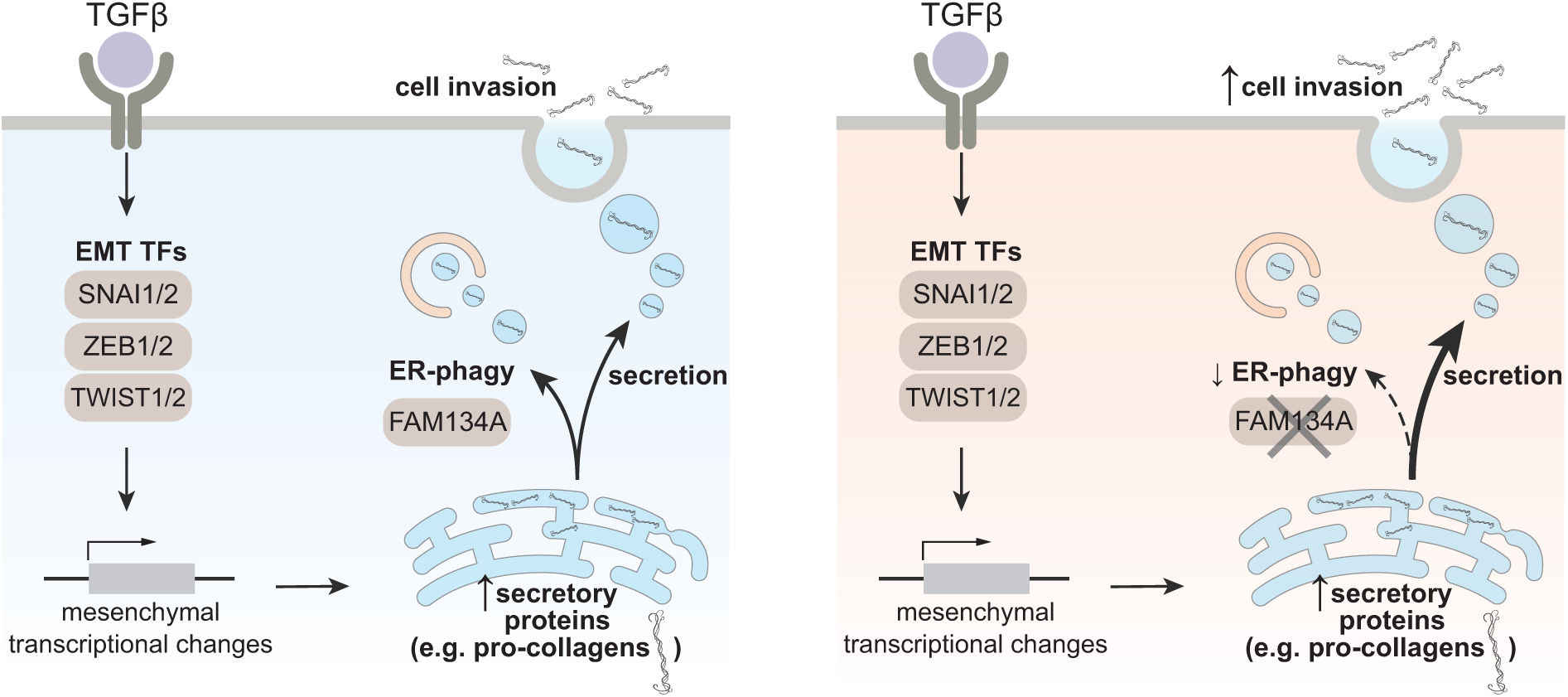
Current working model. Proposed mechanism in which TGFβ-induced upregulation of secretory protein synthesis in the ER is coupled with FAM134A-dependent ER-phagy to fine-tune protein secretion. Loss of FAM134A reroutes procollagens from ER-phagy to exocytosis which promotes cell invasion.

Consistent with this, higher expression of FAM134A correlates with better survival outcome in KIRC datasets, suggesting that FAM134A may act as a tumour suppressor that regulates the interaction between cancer cells and the tumor microenvironment. Although direct loss of function mutations or deletions of FAM134A in cancer cell lines have not been extensively documented, its expression has been shown to be downregulated via microRNA-mediated silencing. For example, secretion of hsa-miR-940 by prostate and breast cancer cells suppresses FAM134A expression and promotes the osteogenic differentiation of human mesenchymal stem cells in vitro ^50^.

EMT remains a major challenge in oncology, driving both metastasis and therapeutic resistance. Targeting EMT and its associated pathways, including ER-phagy, offers a promising avenue for limiting cancer progression and improving treatment outcomes. By characterising FAM134A as a key regulator that controls secretion and invasion, our work demonstrates a potential therapeutic possibility linking ER quality control to cancer progression. These findings lay the foundation for future investigations into FAM134A as a biomarker of metastatic potential, and support the exploration of ER-phagy activation, either through restoring or enhancing FAM134A expression or activity, as a potential therapeutic strategy to limit EMT-driven cancer metastasis.

## Materials and Methods

### Plasmid cloning

All open reading frames (ORFs) are cloned using Gibson Assembly into the pLenti-X1-Neo backbone (digested using BamHI and XbaI restriction enzymes). ORFs are PCR amplified from cDNA reverse-transcribed from HCT116 or A549 mRNA. For BromoTag-FAM134A (wild-type and LIR mutant), the EF1alpha promoter is replaced with the endogenous promoter of FAM134A (950bp upstream of translation start site) to ensure near-physiological expression levels. All CRISPRi and CRISPRa sgRNA constructs are cloned using annealing ligation method into pCRISPRia-v2 (Addgene #84832) following their standard method in Horlbeck et al.^14^. To generate dual/triple sgRNA targeting plasmids, we used Gibson assembly, to incorporate human, mouse and bovine U6 promoters and the respective sgRNA sequences. All of the protospacer sequences used in this study are shown in **Supplementary Table 1.**

To generate knock-ins of mCherry-COL4A1 and mCherry-eGFP-COL4A1, we cloned the sgRNA sequence (COL4A1: CCGGGAACTCACCTTCGCAG) into pX458 (Addgene #48138) using the same annealing ligation approach. The repair template for sp-mCherry-COL4A1 and sp-mCherry-eGFP-COL4A1 were assembled using Gibson assembly by integrating mCherry between the upstream 928bp homology region and downstream 916bp homology region. Note that we inserted the fluorescent tag after the endogenous signalling peptide sequence of COL4A1 (amino acid sequence: MGPRLSVWLLLLPAALLLHEEHSRAAA) to ensure correct targeting of COL4A1 to the ER. To generate FAM134A gene knockout (TCTGAGCCTAGGCATGAGTG), the protospacer sequence was cloned into pX459 (Addgene #48139) using the same annealing ligation approach. All plasmids used in this study are listed in **Supplementary Table 2**.

**Supplementary Table 1.**
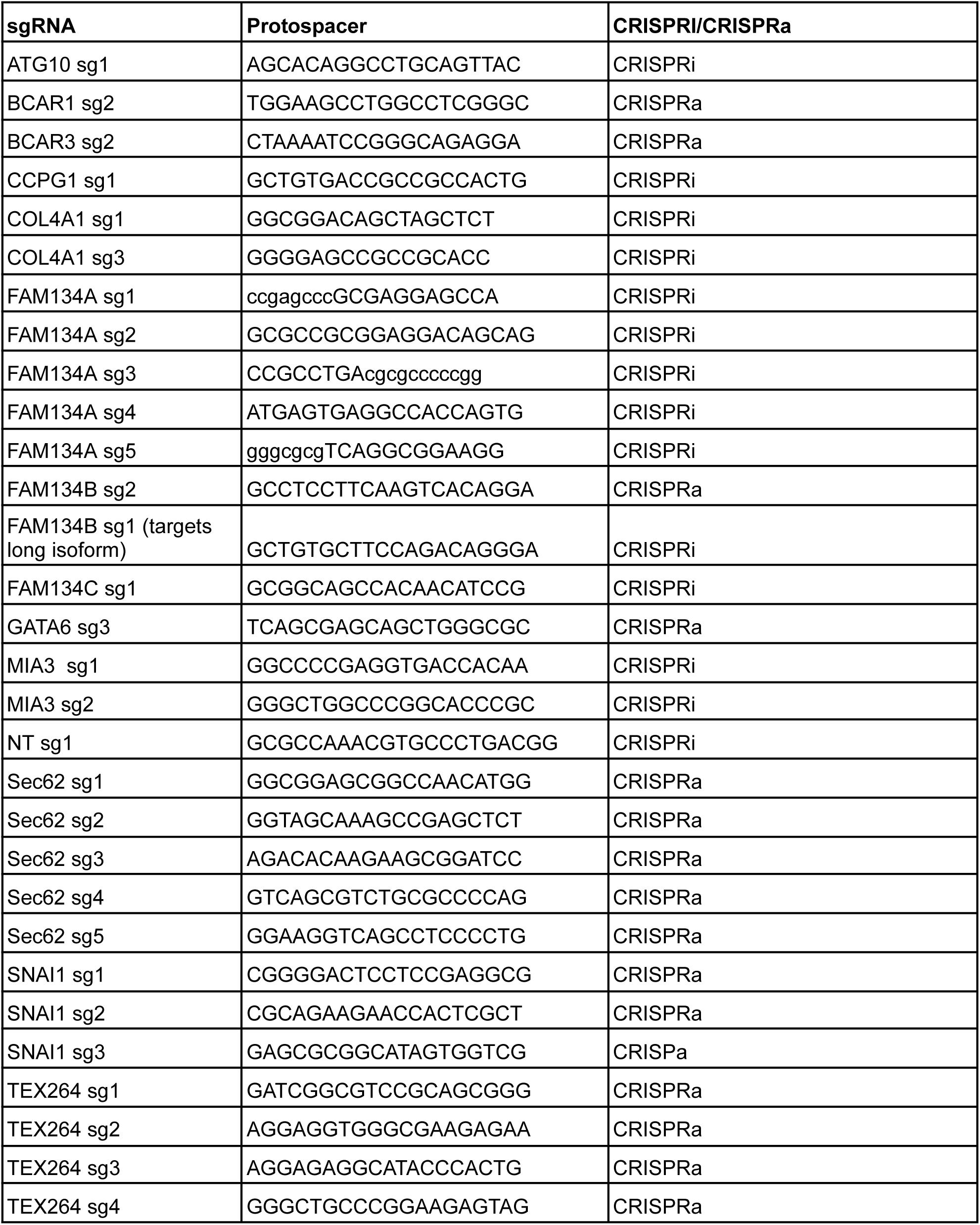
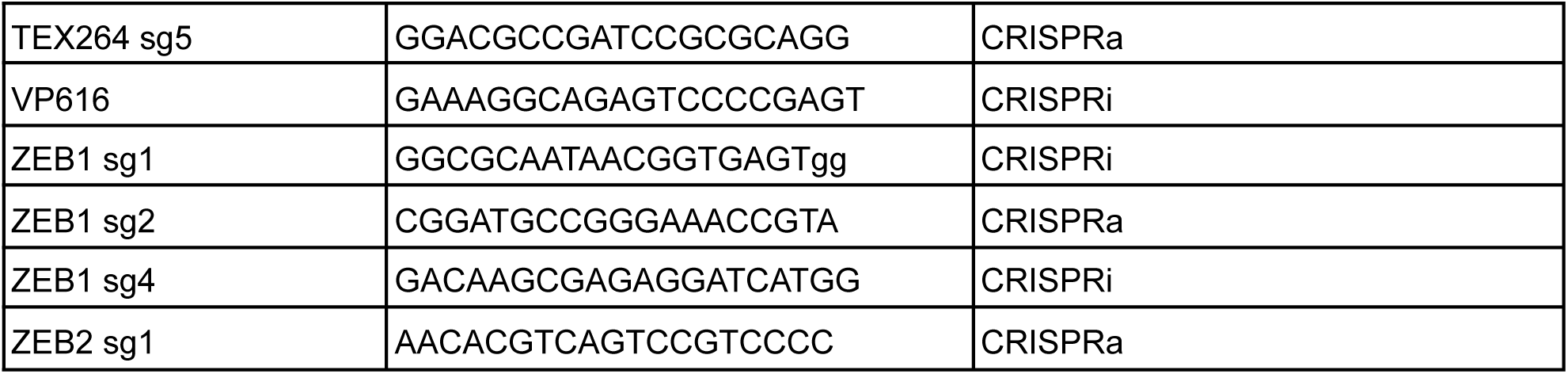
CRISPR single-guide target sequences.

**Supplementary Table 2.** List of plasmids.

|  |  |
| --- | --- |
| pHR-EF1a-dCas9-HA-BFP-KRAB-NLS | Liang et al. 2018 )(Addgene #102244) |
| dCas9-VPR-BFP | This study |
| pGL1-puro-Non-targeting sg1 | Liang et al. 2018 (Addgene #109002) |
| pGL1-Neo-Non-targeting sg1 | This study |
| pGL1-puro-FAM134B-sgA-2 | This study |
| pGL1-puro-TEX264-sgA-1 | This study |
| pGL1-puro-CCPG1-sgA1 | This study |
| pGL1-puro-ZEB1-sgA2 | This study |
| pGL1-puro-SNAI1-sgA1 | This study |
| pGL1-puro-ZEB2-sgA1 | This study |
| pGL1-puro-GATA6-sgA2 | This study |
| pGL1-puro-BCAR1-sgA2 | This study |
| pGL1-puro-BCAR3-sgA2 | This study |
| pLenti-X1-Neo-HA-SNAI1 | This study |
| pLenti-X1-Neo-HA-ZEB1 | This study |
| pLenti-X1-Neo-HA-mTWIST1 | This study |
| pLenti-X1-Neo-HA-GATA6 | This study |
| pLenti-X1-Neo-HA-OVOL1 | This study |
| pGL1-puro-ZEB1-sgi-1 | This study |
| pGL1-puro-ZEB1-sgi-4 | This study |
| pGL1-puro-TSS-GWV2-ATG10-sg1 | Liang et al. 2018 |
| pGL1-puro-TSS-GWV2-sgVPS16 | Liang et al. 2018 |
| pX458-sgCOL4A1 | This study |
| pUC19-sigpep-mCherry-COL4A1 HDR template | This study |
| pUC19-spmCherry-eGFP-COL4A1-HDR template | This study |
| pLenti-X1-BLAST-TMEM192-GFP | This study |
| pGL1-puro-TSS-GWV2-sgFAM134A-sg2 | This study |
| pGL1-puro-FAM134B-iso1-sgi1 | This study |
| pGL1-puro-TSS-GWV2-sgFAM134C-sg1 | This study |
| pGL1-puro-FAM134A-sgi2-FAM134Biso1-sgi1 | This study |
| pGL1-Neo-FAM134B-iso1-sgi1-FAM134C-sgi2 | This study |
| pGL1-puro-triple-FAM134A-sgi2-FAM134Biso1-sgi1-FAM134C-sg | This study |
| pLenti-X1-blast-mCherry-FAM134A-WT | This study |
| pLenti-X1-blast-mCherry-FAM134A-LIRMut | This study |
| pLenti-X1-Neo-mCherry-FAM134B-001 | This study |
| pLenti-X1-Neo-mCherry-FAM134B-iso1-LIRMut | This study |
| pLenti-X1-blast-mCherry-FAM134C-WT | This study |
| pLenti-X1-blast-mCherry-FAM134C-LIRMut | This study |
| pLenti-X1-Neo-HA-FAM134A-WT | This study |
| pLenti-X1-Neo-HA-FAM134A-LIR | This study |
| pLenti-X1-Neo-HA-FAM134B-WT | This study |
| pLenti-X1-Neo-HA-FAM134B-LIR | This study |
| pLenti-X1-Neo-HA-FAM134C-WT | This study |
| pLenti-X1-Neo-HA-FAM134C-LIR | This study |
| pLenti-X1-Neo-endoprom-BromoTag-FAM134A-WT | This study |
| pLenti-X1-Neo-endoprom-BromoTag-FAM134A-LIRmut | This study |
| pLG1-neo-MIA3-sgi1<br>[No BFP] | This study |
| pLG1-neo-MIA3-sgi2<br>[No BFP] | This study |
| pLenti-X1-Neo-HA-FAM134A-RHD-FAM134Biso1-IDR-LIRmut_V | This study |
| pLenti-X1-Neo-HA-FAM134A-RHD-FAM134C-IDR-LIRmut_V2 | This study |
| pLenti-X1-Neo-HA-FAM134B-RHD-FAM134A-IDR-LIRmut_V2 | This study |
| pLenti-X1-Neo-HA-FAM134C-RHD-FAM134A-IDR-LIRmut_V2 | This study |

#### Cell Culture

All cell lines were cultured at 37 °C with 5% CO₂ in a humidified atmosphere. A549, HCT116, DLD1, U2OS, MCF7, HeLa, HEK293T and NMuMG cells were cultured in DMEM medium supplemented with 10% FBS, 0.1 mM non-essential amino acids (GIBCO), 1 mM sodium pyruvate (GIBCO), 100 U/mL penicillin (GIBCO),and 100 g/mL streptomycin (GIBCO). SH-SY5Y cells were cultured in DMEM/F12 media with the same supplements. All cell lines were routinely tested to be mycoplasma free.

#### Cell Treatments

ER-phagy was induced with media starvation using EBSS with calcium, magnesium, and phenol red (GIBCO; Cat. No.: 24010043). For EATR-FACS assay, cells were plated 48 hr prior to EBSS treatment. EATR expression is induced using 1µg/ml doxycycline 24 hr prior to amino acid starvation. Starvation treatment was carried out for 16 hr. Cells in fed conditions indicate incubation in complete DMEM described above.

Unless stated otherwise, cells were treated with 5ng/µL TGFβ-1 (Proteintech HZ-1011) for 48 hr. SB505124 (1µM) was co-administered at the time of TGFβ treatment. Concanamycin (50nM or 100nM) and protease inhibitor cocktail (Pepstatin A- 5µM; Leupeptin- 10µM, E64D - 10µM) were administered to cells 24 hr after TGFβ treatment concentration. To inhibit MEK1/2, cells were treated with Trametinib at a concentration of 10 mM and 100nM (Cambridge Biosciences, T5857) for 16 hr along with EBSS.

#### Lentiviral packaging and transduction

Lentiviral packaging was performed in HEK293T cells using Opti-MEM (GIBCO; Cat. No.: 31985062). Briefly, delta-VPR (Addgene #8455), VSVG (Addgene #8454), and the construct of interest was incubated at the ratio of 4:1:5 in Opti-MEM and PEI for 10min. The mixture is then added dropwise to HEK293T cells. Lentiviral supernatant was harvested at 48hr post-transfection and HEK293T cells were replenished with fresh media for another harvest at 72 hr post-transfection. Lentiviral supernatant was added to the culturing cells for 24 hr before replacing with fresh media. Antibiotic selection was initiated 48hr post transduction.

#### RNA interference

A549 cells were transfected with Human FAM134A, FAM134B, FAM134C and negative control siRNA (Dharmacon) at a concentration of 50 nM using Lipofectamine RNAiMAX (Thermo Fisher Scientific; Cat. No.: 13778100) according to the manufacturer’s instructions. Briefly, siRNA (50 µM; 2 µL) was added into 250 µL Opti-MEM and incubated for 5 min. Subsequently, 3 µL RNAiMAX was diluted in 250 µL Opti-MEM was mixed into the siRNA solution and incubated for 20 min. Cells were seeded at around ∼20% confluency and transfected on the same day. This was followed by incubation in transfection media for 48 hr. Cells were then trypsinized, seeded (∼20%) and TGFβ treated for 48 hr. Following this, cells were harvested for downstream applications including Western blotting, flow cytometry and transwell invasion assay. The siRNA sequences are provided in **Supplementary Table 3.**

**Supplementary table 3.**
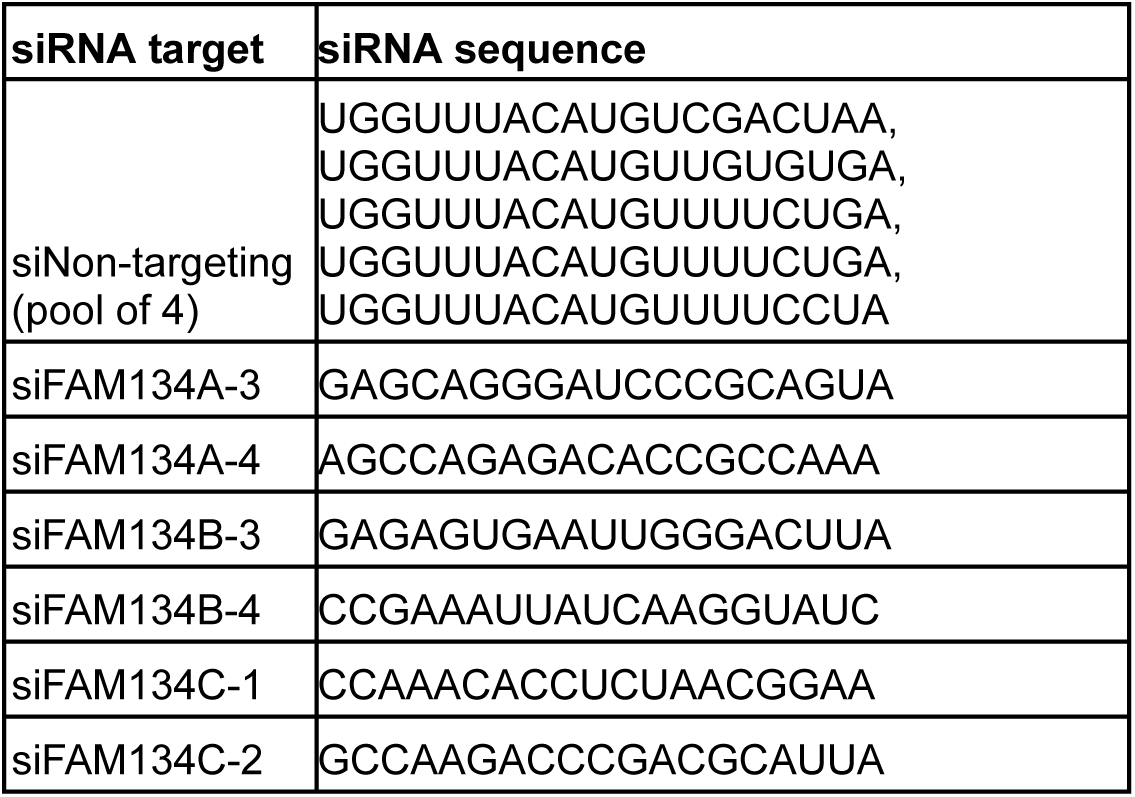
List of siRNA sequences.

#### CRISPRi and CRISPRa cell line generation

HCT116 and A549 cells were transduced with dCas9-KRAB-BFP (CRISPRi; Addgene# 102244) or dCas9-VPR-BFP (CRISPRa). A week post-transduction, the cells were single cell-sorted via FACS based on BFP expression. CRISPRi and CRISPRa clones were then transduced with control sgRNAs to functionally validate their knockdown and overexpression efficiency using Western blotting. CRISPR sequences used in this study are tabulated in **Supplementary Table 1.**

#### CRISPR/Cas9-mediated knock-out or knock-in of fluorescence tags

A549 cells were seeded a day before such that the cells are 70-80% confluent on the day of transfection. For knock-out of FAM134A, 1µg of pX459-sgFAM134A (TCTGAGCCTAGGCATGAGTG) were mixed with 2µL of P3000 in 75µL of Opti-MEM. In parallel, 3µL of Lipofectamine 3000 was mixed with 75µL of Opti-MEM. The two mixtures were combined and incubated at room temperature for 10 min before being added dropwise into the cells.

For knock-in of mCherry/ mCherry-eGFP tag to COL4A1 loci, 0.5µg of pX459-sgCOL4A1 (protospacer seq: CCGGGAACTCACCTTCGCAG) and 1µg of pUC19-sp-mCherry-eGFP-COL4A1 HDR template (or pUC19-sp-mCherry-COL4A1 HDR template) were mixed with 3µL of P3000 reagent in 75µL of OPTIMEM. In parallel, 4.5µL of Lipofectamine 3000 was mixed with 75µL of OptiMEM. The two mixtures were then combined and incubated at room temperature for 10 min before being added dropwise into the cells. Cells were replaced with fresh media 24 hr post-transfection.

Puromycin was added to the transfected cells 48 hr post-transfection (2µg/mL, for 48 hr). Cells were then left in complete media (without puromycin) for 1-2 weeks until positive cells recovered to full confluency. Puromycin selection was sufficient to generate a pool of near-complete FAM134A knockout without a need for single cell clone isolation. To isolate successful mCherry/mCherry-eGFP-COL4A1 knock-in clones, cells were treated with TGFβ for 48 hr to induce collagen expression and mCherry positive clones were sorted by FACS into 96-well plates.

#### Genome-wide CRISPRa screen

The sgRNA sequences for genome-wide screening were based on the Weissman CRISPRa-v2 pooled library (Addgene #83978) and contained 5 sgRNAs per gene. The entire genome-wide CRISPRa screen described below was performed at a minimum of 500-fold coverage, which equals approximately 52 millions cells for each step. HCT116-dCas9-VPR-ssCALR-mCherry-eGFP-KDEL cells were transduced with the human CRISPRa-V2-TOP5 sgRNA lentiviral library from the Weissman lab (Addgene #83978) at 1:4 lentiviral dilution (based pre-optimised dilution ratio to achieve 15-20% BFP+ve cells). After 48 hr of transduction, cells were treated with 4µg/ml of puromycin for 5 days to achieve >95% BFP positive cells.

For each biological replicate, 52 million cells were trypsinized and collected as unsorted background reference samples. For FACS, 150 million cells were collected and passed through a mesh strainer to remove cell clumps. The high cell number compensated for cell death from amino acid starvation and non-single clumps that were omitted from analysis. The ‘enhanced’ and ‘inhibited’ FACS gates were set up based on the mCherry and eGFP signals such that 25% each of the top and bottom cell distribution were collected (i.e. 13 million cells in each gate). The entire FACS was performed using either untreated cells, or amino-acid starved (EBSS; 16 hr) cells. In each case, two biological replicates were performed.

These sorted samples, together with the unsorted ‘background’ samples were then subjected to genomic DNA (gDNA) extraction using Puregene Gentra kit (Qiagen) following manufacturer’s protocol. Next, 500µg and 150µg of genomic DNA for background and sorted samples, respectively, were used to perform indexing PCR of the sgRNA cassette using NEBNext Ultra II Q5 Mastermix (NEB; 10µg of gDNA per 100µL of PCR reaction). The amplicons were cleaned up using SeraMag SPRI beads to remove primers and genomic DNA. Amplicon concentrations were determined using Qubit™ 1x dsDNA HS Assay Kits prior to pooling according to the expected sequencing read ratio for each sample. Next generation sequencing (NGS) was performed using NextSeq500 single-end read 100 with 20% PhiX spiked in. The enrichment of sgRNAs in the sorted samples relative to the background samples were then analysed using the Screen Processing pipeline ^16^ ^51^.

#### Western Blotting

Cells were lysed in RIPA buffer (Merck; Cat. No.: 20-188), supplemented with 1x Halt Protease Inhibitor Cocktail (Thermo Fisher Scientific; Cat. No.: 78429), at 4 °C for 15 min and then spun at 20,000g for 10 min to remove insoluble debris. Protein concentrations were quantified by Bradford assay (Thermo Fisher ; Cat. No.: A55866). Lysates were normalised based on protein concentration and 4x NuPage LDS Sample Buffer (Thermo Fisher Scientific; Cat. No.: NP0008) supplemented with B-mercaptoethanol (5% v/v). Samples were boiled at 98 °C for 5 min. For secretion analysis, conditioned media were collected on the same day as cell lysate. The conditioned media were normalised based on the cell lysate concentration determined using Bradford assay.

Between 20-40 µg of samples were run on NuPAGE Bis-Tris 4%–12% gels in NuPage MES SDS Buffer (Invitrogen) for 45 min at 180 V and transferred to 0.4-mm nitrocellulose membranes via wet transfer at 1.3 A and 90 V for 90 min. Membranes were blocked with 5% (w/v) milk in Tris-buffered saline containing 0.1% Tween 20 (TBS-T) for 30 min, and subsequently washed with TBS-T three times. All primary antibodies were diluted at the appropriate concentration in 5% BSA (w/v) in TBS-T except the FAM134B antibody which was diluted in 5% milk (w/v) in TBS-T. The membrane was incubated in the primary antibody for either 4 h at room temperature or overnight at 4 °C. The membrane was washed with TBS-T three times for five min each. The blots were incubated for 30 min in the milk solution with a 1:30,000 dilution of Li-Cor near-infrared fluorescence secondary antibodies. The blots were scanned using Li-Cor’s Near-InfraRed fluorescence Odyssey CLx Imaging System, and densitometry quantifications were done using Li-Cor’s ImageStudio software complementary to Odyssey. Antibodies used in this study are outlined in **Supplementary Table 4**.

**Supplementary Table 4.** Antibodies used in western blot studies (Rb - rabbit; Ms - mouse)

| Antigen name (ID) | Company | Cat No. | Species |
| --- | --- | --- | --- |
| FAM134B | ProteinTech | 21537-1-AP | Rb |
| TEX264 | Sigma | HPA017739 | Rb |
| SEC62 | Abcam | ab140644 | Rb |
| GAPDH (D4C6R) | CST | #97166 | Ms |
| Cas9 (7A9-3A3) | CST | 14697 | Ms |
| E-cadherin (4A2) | CST | 14472 | Ms |
| Vimentin (E-5) | Santa Cruz | sc-373717 | Ms |
| ZEB1 (D80D3) | CST | 3396 | Rb |
| HA (C29F4) | CST | 3724 | Rb |
| B-actin (C4) | Santa Cruz | sc-47778 | Ms |
| mCherry | Abcam | ab183628 | Rb |
| FAM134A/ RETREG2 | Fortis Life Science | A305-801A | Rb |
| FAM134C/ RETREG3 | Invitrogen | PA5-53433 | Rb |
| Calnexin (E-10) | Santa Cruz | SC-46669 | Ms |
| LC3B | Novus Biologicals | NB100-2220 | Rb |
| N-cadherin (D4R1H) | CST | 13116 | Rb |
| ATG10 | Abcam | ab124711 | Rb |
| PDI (H160) | Santa Cruz | sc-20132 | Rb |
| COL4A1 | CST | #50273 | Rb |
| PAI-1 | ProteinTech | 13801-1-AP | Rb |
| BiP | CST | C50B12 | Rb |
| COL5A1 (C-5) | Santa Cruz | sc-166155 | Ms |
| MIA3/TANGO1 | Abcam | ab244506 | Rb |

### Immunofluorescence

Cells were fixed in 4% (wt/vol) paraformaldehyde for 15 min followed by permeabilization and blocking using 0.1% digitonin in 5% BSA for 15 min. LAMP2 anti-mouse (Santa Cruz) was prepared in 0.1% digitonin/5%BSA and incubated for 1hr at room temperature. This was followed by three PBS washes for 5 min each. Alexa Fluor488 anti-rabbit IgG secondary antibody, prepared in 0.1% digitonin/5%BSA, was incubated for 30 min at room temperature, followed by three PBS washes for 5 min each. Coverslips were mounted onto glass slides using ProLong Gold Antifade reagent with or without DAPI addition for nucleus visualisation. Images were taken using a 63x or 100x oil objective lens and post-processed in Adobe Photoshop for specific inset enlargement and RGB channel separation. Colocalisation analysis was carried in Imaris 10 using the Spot Colocalisation plug-in.

#### Flow cytometry for ER-phagy

Flow cytometry was performed using an LSR Fortessa Flow Cytometer and subsequent analysis was performed using FlowJo 10.1. All experiments were performed using live cells to prevent reversal of eGFP quenching. The intensities for both eGFP and mCherry of the EATR cells at fed condition were used as references to define the gate for zero ER-phagy.

For mCherry-eGFP-COL4A1 cells, since a fraction of COL4A1 is constitutively targeted for ER-phagy upon expression, we used 24 hr ConA (50µM) treatment to define the gate for zero ER-phagy. COL4A1 ER-phagy detection is based on the shift of cell population into the ER-phagy gate following 48 hr TGFβ stimulation. For experimental conditions interrogating the effects of gene knock-down conditions, COL4A1 ER-phagy was gated at ∼20% in the non-targetting condition as a reference. For mCherry-COL4A1 analysis, the total percentage of mCherry cells were determined relative to 50% of the total mCherry positive cells in the TGFβ treated control condition. On average, 10,000 live, single cells were analyzed per condition and all statistical analyses were performed using data from at least three biological replicates.

#### qRT-PCR

RNA extraction was performed using the RNAeasy miniprep kit (Qiagen; Cat. No.: 74104) according to the manufacturer’s instructions. 1mg of RNA per sample was used for reverse transcription using iScript™ Reverse Transcription Supermix for RT-qPCR (Bio-Rad; Cat. No.: 1708840) according to the manufacturer’s instructions. qRT-PCR reactions were set up using SsoFast EvaGreen Supermix with low ROX (Bio-Rad; Cat. No.: 1725211) and run in triplicates using StepOnePlus Real-Time PCR system (Applied Biosystems) according to manufacturer’s instructions. A complete list of all primers used are compiled in **Supplementary Table 5.**

**Supplementary Table 5.**
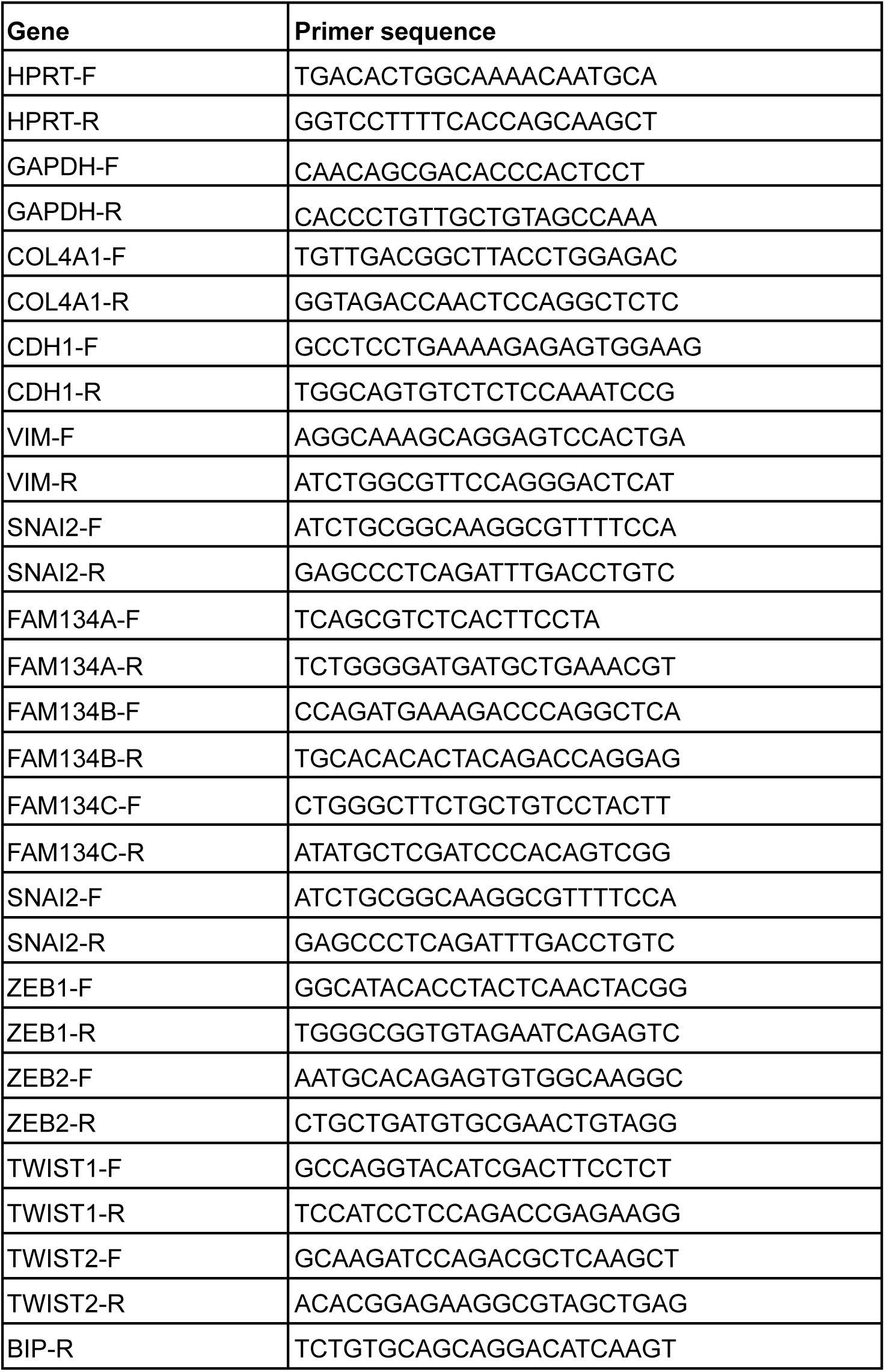
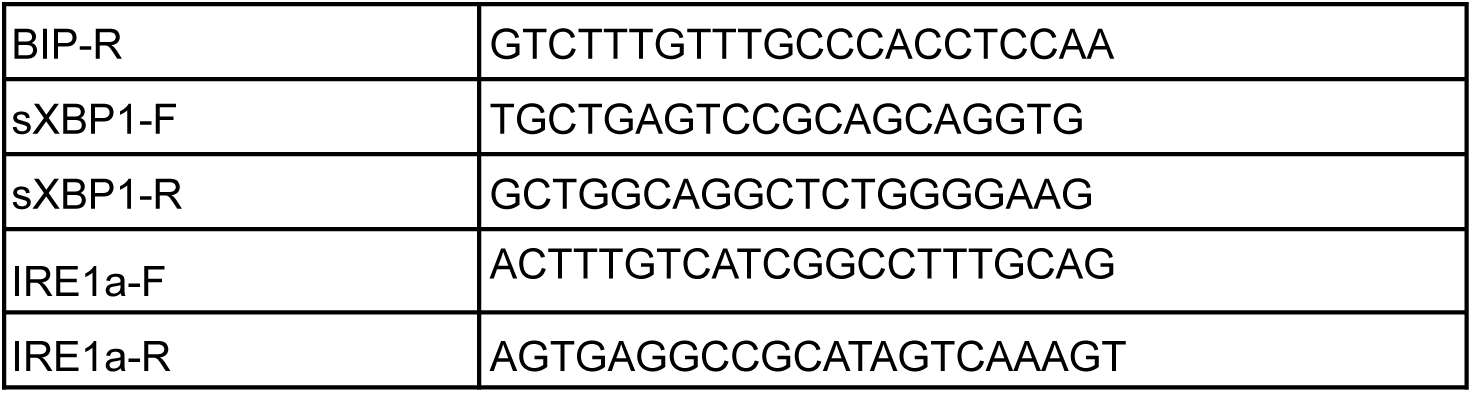
List of qRT-PCR primers.

#### Whole cell lysate mass spectrometry

A549 cells were seeded in 6-well plates at a density of 250,000 cells/well, and left to attach for 24 hr before treatment with TGFβ (5ng/ml) for 48h and ConA (50 nM) for the last 16 hr of TGFβ treatment. Cells were harvested by trypsinisation, washed in PBS, and lysed in lysis buffer (5% SDS (Sigma Aldrich, Cat. No.: 436143), 50 mM TEAB (Sigma Aldrich, Cat. No.: T7408), pH 8.5). Samples were sonicated on a Diagenode Bioruptor® Plus in high power mode for 30 cycles (sonication cycle: 30 seconds ON, 30 seconds OFF) to shear genomic DNA, and protein content was quantified by BCA assay (Thermo Fisher Scientifc, Cat. No.: A55864). Samples were normalised to 100 µg of protein in 46 µL of lysis buffer and prepared using Protifi S-TRAP Mini (Cat. No.: C002-MINIX-0010/40/80PK) following the manufacturer’s protocol. Briefly, disulfide bonds were reduced with the addition of 2 μL of 125 mM TCEP (Thermo Fisher Scientific, Cat. No.: 77720) and incubation at 55 °C for 15 min, then alkylated with the addition of 2 μL of 530 mM MMTS (Thermo Fisher Scientific, Cat. No.: 23011) and incubation at room temperature for 10 min. Finally, 5 μL of 12% phosphoric acid and 350 μL of binding/wash solution (100 mM TEAB in 90% methanol) were added to the samples. Proteins were bound to S-TRAP Mini columns and washed four times with 400 μL of binding/wash solution. On-column digestion was performed by addition of 20 µg Pierce mass spectrometry grade Trypsin-protease (Thermo Fisher Scientific, Cat. No.: 90057) diluted in 125 µL 50 mM TEAB and incubation at 37 °C overnight. Digested peptides were eluted sequentially in single 80 µL washes of each 50 mM TEAB, 0.2% (v/v) formic acid (Fisher Scientific, Cat. No.: A117-50), and 50% (v/v) acetonitrile. Solutions were evaporated to dryness using a Savant SpeedVac SPD140DDA vacuum concentrator connected to a Savant RVT5105 refrigerated vapor trap.

Peptides were resuspended in 200 μL 0.1% formic acid with 0.015% DDM and analysed by injecting 0.5 μL onto a Vanquish Neo UHPLC System coupled with an Orbitrap Astral Mass Spectrometer (Thermo Fisher Scientific). Peptides were loaded onto a PepMap Neo Trap Cartridge (C18, 5 µm, 5 mm × 300 µm; Thermo Fisher Scientific, Cat. No.: 174500) and analysed on a C18 EASY-Spray HPLC Column (C18, 2 µm, 150 mm ×150 μM; Thermo Fisher Scientific, Cat. No.: ES906) with an 11.8 minute gradient from 1% to 55% Buffer B (Buffer A: 0.1% formic acid in water; Buffer B: 0.08% formic acid in 80:20 acetonitrile:water). The gradient was established as follows: 0.7 min at 1.8 µL/min from 1% to 4% B, 0.3 min at 1.8 µL/min from 4% to 8% B, 6.7 min at 1.8 µL/min from 8% to 22.5% B, 3.7 min at 1.8 µL/min from 22.5% to 35% B, 0.4 min at 2.5 µL/min from 35% to 55% B. Eluted peptides were analysed using data-independent acquisition (DIA) mode on the mass spectrometer by the MRC PPU Mass Spectrometry facility, University of Dundee.

DIA data were processed in DIA-NN (v2.0 Academia) using a library-free workflow with in silico spectral library prediction. Searches were conducted against a custom human UniProt/Swiss-Prot database including isoforms and cRAP contaminants. Trypsin specificity was used with one missed cleavage allowed. Peptides (7–30 aa; m/z 300–1800; charges +1–4) were considered. Methylthio modification of cysteine (UniMod:39) was set as fixed, N-terminal methionine excision was enabled, and identifications were filtered at 1% FDR.

For downstream analysis, protein Groups (PGs) identified with a single peptide were removed. PGs with less than 75% valid values in at least one condition were also removed. Raw intensities were log2 transformed and median normalised (by subtraction of sample median followed by addition of global/all samples median). Missing values were imputed using random draws from a normal distribution, with the mean being the value corresponding to the 1st percentile of the sample distribution, and standard deviation being sample median (only considering PGs present with more than 50% non-missing values). Differentially expressed proteins were identified using a moderated t-test performed using the Limma statistical package. Analysis was carried out by MRC PPU Mass Spectrometry facility, University of Dundee.

#### FAM134 RHD and IDR Domain Swapping

The FAM134 paralogues have four transmembrane helixes followed by a conserved amphipathic helix. To generate the FAM134 RHD-IDR hybrids, we define the RHD/IDR border for each paralogue at the last amino acid of the conserved amphipathic helix (see Supp Fig 6B). The respective RHD and IDR (LIR-mutant version) fragments are PCR-amplified and Gibson assembled into a pLenti-X1-Neo construct cut at BamHI/XbaI sites.

#### Survival Analysis using GEPIA2

To assess the prognostic significance of FAM134A, FAM134B, and FAM134C expression in human cancers, we used GEPIA2 (http://gepia2.cancer-pku.cn), an interactive web platform based on TCGA and GTEx RNA-seq datasets. Kaplan–Meier survival analysis was conducted using the "Survival Analysis" module of GEPIA2 against either Pan-Cancer dataset and Kidney Renal Clear Cancer Cell Carcinoma (KIRC). Each gene was queried individually and we selected Overall Survival (OS) as the clinical endpoint. The quartile cutoff option was chosen to define high and low expression groups, where patients in the top 25% and bottom 25% of gene expression levels were compared. The log-rank test was used to assess statistical significance, and the hazard ratio (HR) with a 95% confidence interval was reported for each analysis. All resulting survival plots were downloaded directly from the GEPIA2 interface. The results generated are in whole or part based upon data generated by the TCGA Research Network: https://www.cancer.gov/tcga.

#### Wound Healing Assay

Culture insert 2-well systems (Ibidi, Cat. No.: 80209) were placed in a 24-well plate using a sterile tweezer. 70 μL of complete media was added into each well prior to cell seeding and incubated for 30 min. After removing the media, 70 μL of cell suspension (8×10^4^ cells/cm^2^ seeding density) was added into each well and incubated for 24 hr at 37 °C to allow cell attachment. Prior to migration assay, cells were starved treating DMEM media including 0% FBS for 3 h. After the incubation, the inserts were gently removed by grabbing one corner using a sterile tweezer, and the cells were washed using PBS to remove nonadherent cells and/or cell debris. Then, the cells were treated with complete media only (control) or media containing 5 ng/ml TGFβ. The gap between cells was imaged using EVOS M5000 imaging system with a 10x objective. The images were analysed by using ImageJ software using the ‘Wound Healing’ plug-in.

### Transwell invasion Assay

Matrigel matrix (Corning) was diluted in serum-free media to a final concentration of 200 µg/mL. 100 µL of diluted Matrigel matrix was added to the center of each Transwell insert (8uM PET membrane, Corning 3464). Inserts were placed in 24-well plates containing 600µL of 10% FBS supplemented media. The plate was incubated at 37 °C for 1 hour to allow the Matrigel matrix to form a gel. Next 200 µL of cells were seeded on top of the Matrigel coated transwells at a density of 9×10^4^/mL, in serum-free media. TGFβ was added to the appropriate cells at a concentration of 5ng/mL and incubated for 48 hr. For conditioned media assays, media containing 1% FBS was collected from siNT and siFAM134A-A549 cells, which has been treated with TGFβ for 48 hr. The conditioned media was plated on A549 parental cells inside Boyden chambers coated with Matrigel. Again, TGFβ was added to the appropriate cells at a concentration of 5ng/mL and incubated for 48 hr. Media was removed from the top of the inserts and the bottom of wells and replaced with 4% (w/v) paraformaldehyde for 10 min. Following fixation, DAPI (1µg/mL in PBS) was added to both the bottom and the top of the transwell plate for 10 min. Transwells were then washed once with PBS. Images were taken on an EVOS M5000 imaging system using a 4x objective. Cells were removed from the top of the transwell by gently scraping with a cotton bud. Imaging was repeated to capture the cells at the bottom of the transwell. Nuclei were counted using ImageJ and invasion index was expressed as a percentage of cells at the bottom of the transwells relative to the top.

#### Statistical Analysis

All statistical analyses were performed using data from three or more independent experiments. All error bars represent standard errors. Depending on the context of the experiment, statistical scores were derived using either Student’s *t* test (paired or unpaired) or ANOVA (with Tukey’s, Holm-Šidák or Dunnett’s multiple comparison test) as stated in each figure legend. Data distribution was assumed to be normal, but this was not formally tested.

## Data Availability

The genome-wide CRISPRa screen data generated in this study have been deposited in the Gene Expression Omnibus (GEO) under the accession number GSE315931. The mass spectrometry proteomics data have been deposited to the ProteomeXchange Consortium via the PRIDE ^52^ ^53^ partner repository with the dataset identifier PXD073014.

## Declaration of generative AI and AI-assisted technologies in the manuscript preparation process

During the preparation of this work, the authors used ChatGPT for spell check and to improve overall clarity of the manuscript. After using this tool, the authors have reviewed and edited the content as needed and thus take full responsibility for the content of the published article.

## Bibliography

1. Schwarz, D.S., and Blower, M.D. (2016). The endoplasmic reticulum: structure, function and response to cellular signaling. Cell. Mol. Life Sci. 73, 79–94. 10.1007/s00018-015-2052-6.

2. Ferro-Novick, S., Reggiori, F., and Brodsky, J.L. (2021). ER-Phagy, ER Homeostasis, and ER Quality Control: Implications for Disease. Trends Biochem. Sci. 46, 630–639. 10.1016/j.tibs.2020.12.013.

3. Fumagalli, F., Noack, J., Bergmann, T.J., Cebollero, E., Pisoni, G.B., Fasana, E., Fregno, I., Galli, C., Loi, M., Soldà, T., et al. (2016). Translocon component Sec62 acts in endoplasmic reticulum turnover during stress recovery. Nat. Cell Biol. 18, 1173–1184. 10.1038/ncb3423.

4. An, H., Ordureau, A., Paulo, J.A., Shoemaker, C.J., Denic, V., and Harper, J.W. (2019). TEX264 Is an Endoplasmic Reticulum-Resident ATG8-Interacting Protein Critical for ER Remodeling during Nutrient Stress. Mol. Cell 74, 891-908.e10. 10.1016/j.molcel.2019.03.034.

5. Chino, H., Hatta, T., Natsume, T., and Mizushima, N. (2019). Intrinsically Disordered Protein TEX264 Mediates ER-phagy. Mol. Cell 74, 909–921.e6. 10.1016/j.molcel.2019.03.033.

6. Liang, J.R., Lingeman, E., Luong, T., Ahmed, S., Muhar, M., Nguyen, T., Olzmann, J.A., and Corn, J.E. (2020). A Genome-wide ER-phagy Screen Highlights Key Roles of Mitochondrial Metabolism and ER-Resident UFMylation. Cell 180, 1160–1177.e20. 10.1016/j.cell.2020.02.017.

7. Stephani, M., Picchianti, L., Gajic, A., Beveridge, R., Skarwan, E., Sanchez de Medina Hernandez, V., Mohseni, A., Clavel, M., Zeng, Y., Naumann, C., et al. (2020). A cross-kingdom conserved ER-phagy receptor maintains endoplasmic reticulum homeostasis during stress. eLife 9. 10.7554/eLife.58396.

8. Nthiga, T.M., Kumar Shrestha, B., Sjøttem, E., Bruun, J.-A., Bowitz Larsen, K., Bhujabal, Z., Lamark, T., and Johansen, T. (2020). CALCOCO1 acts with VAMP-associated proteins to mediate ER-phagy. EMBO J. 39, e103649. 10.15252/embj.2019103649.

9. Khaminets, A., Heinrich, T., Mari, M., Grumati, P., Huebner, A.K., Akutsu, M., Liebmann, L., Stolz, A., Nietzsche, S., Koch, N., et al. (2015). Regulation of endoplasmic reticulum turnover by selective autophagy. Nature 522, 354–358. 10.1038/nature14498.

10. Grumati, P., Morozzi, G., Hölper, S., Mari, M., Harwardt, M.-L.I.E., Yan, R., Müller, S., Reggiori, F., Heilemann, M., and Dikic, I. (2017). Full length RTN3 regulates turnover of tubular endoplasmic reticulum via selective autophagy. eLife 6. 10.7554/eLife.25555.

11. Smith, M.D., Harley, M.E., Kemp, A.J., Wills, J., Lee, M., Arends, M., von Kriegsheim, A., Behrends, C., and Wilkinson, S. (2018). CCPG1 Is a Non-canonical Autophagy Cargo Receptor Essential for ER-Phagy and Pancreatic ER Proteostasis. Dev. Cell 44, 217–232.e11. 10.1016/j.devcel.2017.11.024.

12. Liang, J.R., Lingeman, E., Luong, T., Ahmed, S., Nguyen, T., Olzmann, J., and Corn, J.E. (2019). A genome-wide screen for ER autophagy highlights key roles of mitochondrial metabolism and ER-resident UFMylation. BioRxiv. 10.1101/561001.

13. Chaffer, C.L., San Juan, B.P., Lim, E., and Weinberg, R.A. (2016). EMT, cell plasticity and metastasis. Cancer Metastasis Rev. 35, 645–654. 10.1007/s10555-016-9648-7.

14. Konermann, S., Brigham, M.D., Trevino, A.E., Joung, J., Abudayyeh, O.O., Barcena, C., Hsu, P.D., Habib, N., Gootenberg, J.S., Nishimasu, H., et al. (2015). Genome-scale transcriptional activation by an engineered CRISPR-Cas9 complex. Nature 517, 583–588. 10.1038/nature14136.

15. Liang, J.R., Lingeman, E., Ahmed, S., and Corn, J.E. (2018). Atlastins remodel the endoplasmic reticulum for selective autophagy. J. Cell Biol. 217, 3354–3367. 10.1083/jcb.201804185.

16. Horlbeck, M.A., Gilbert, L.A., Villalta, J.E., Adamson, B., Pak, R.A., Chen, Y., Fields, A.P., Park, C.Y., Corn, J.E., Kampmann, M., et al. (2016). Compact and highly active next-generation libraries for CRISPR-mediated gene repression and activation. eLife 5. 10.7554/eLife.19760.

17. Abe, H., Kikuchi, S., Hayakawa, K., Iida, T., Nagahashi, N., Maeda, K., Sakamoto, J., Matsumoto, N., Miura, T., Matsumura, K., et al. (2011). Discovery of a Highly Potent and Selective MEK Inhibitor: GSK1120212 (JTP-74057 DMSO Solvate). ACS Med. Chem. Lett. 2, 320–324. 10.1021/ml200004g.

18. Salomó Coll, C., Di Monaco, M., Holkham, J., Smith, M., Muir, M., Gautier, P., Dunn-Davies, H., Zheng, X., Krishnankutty, R., Kemp, A.J., et al. (2025). ER-phagy and proteostasis defects prime pancreatic epithelial state changes in KRAS-mediated oncogenesis. Dev. Cell. 10.1016/j.devcel.2025.07.016.

19. Nieto, M.A. (2002). The snail superfamily of zinc-finger transcription factors. Nat. Rev. Mol. Cell Biol. 3, 155–166. 10.1038/nrm757.

20. Spaderna, S., Schmalhofer, O., Wahlbuhl, M., Dimmler, A., Bauer, K., Sultan, A., Hlubek, F., Jung, A., Strand, D., Eger, A., et al. (2008). The transcriptional repressor ZEB1 promotes metastasis and loss of cell polarity in cancer. Cancer Res. 68, 537–544. 10.1158/0008-5472.CAN-07-5682.

21. Ichikawa, M.K., Endo, K., Itoh, Y., Osada, A.H., Kimura, Y., Ueki, K., Yoshizawa, K., Miyazawa, K., and Saitoh, M. (2022). Ets family proteins regulate the EMT transcription factors Snail and ZEB in cancer cells. FEBS Open Bio 12, 1353–1364. 10.1002/2211-5463.13415.

22. Fan, C., Wang, Q., van der Zon, G., Ren, J., Agaser, C., Slieker, R.C., Iyengar, P.V., Mei, H., and Ten Dijke, P. (2022). OVOL1 inhibits breast cancer cell invasion by enhancing the degradation of TGF-β type I receptor. Signal Transduct. Target. Ther. 7, 126. 10.1038/s41392-022-00944-w.

23. Frisch, S.M., Farris, J.C., and Pifer, P.M. (2017). Roles of Grainyhead-like transcription factors in cancer. Oncogene 36, 6067–6073. 10.1038/onc.2017.178.

24. Liu, H., Du, F., Sun, L., Wu, Q., Wu, J., Tong, M., Wang, X., Wang, Q., Cao, T., Gao, X., et al. (2019). GATA6 suppresses migration and metastasis by regulating the miR-520b/CREB1 axis in gastric cancer. Cell Death Dis. 10, 35. 10.1038/s41419-018-1270-x.

25. Guaita, S., Puig, I., Franci, C., Garrido, M., Dominguez, D., Batlle, E., Sancho, E., Dedhar, S., De Herreros, A.G., and Baulida, J. (2002). Snail induction of epithelial to mesenchymal transition in tumor cells is accompanied by MUC1 repression and ZEB1 expression. J. Biol. Chem. 277, 39209–39216. 10.1074/jbc.M206400200.

26. Miyafuji, Y., Zhong, X., Uchida, I., Koi, M., and Hemmi, H. (2001). Growth inhibition due to complementation of transforming growth factor-beta receptor type II-defect by human chromosome 3 transfer in human colorectal carcinoma cells. J. Cell. Physiol. 187, 356–364. 10.1002/jcp.1084.

27. Lee, J., Ballikaya, S., Schönig, K., Ball, C.R., Glimm, H., Kopitz, J., and Gebert, J. (2013). Transforming growth factor beta receptor 2 (TGFBR2) changes sialylation in the microsatellite unstable (MSI) Colorectal cancer cell line HCT116. PLoS ONE 8, e57074. 10.1371/journal.pone.0057074.

28. DaCosta Byfield, S., Major, C., Laping, N.J., and Roberts, A.B. (2004). SB-505124 is a selective inhibitor of transforming growth factor-beta type I receptors ALK4, ALK5, and ALK7. Mol. Pharmacol. 65, 744–752. 10.1124/mol.65.3.744.

29. Myllyharju, J. (2003). Prolyl 4-hydroxylases, the key enzymes of collagen biosynthesis. Matrix Biol. 22, 15–24. 10.1016/S0945-053X(03)00006-4.

30. The effect of transforming growth factor-beta on cell proliferation and collagen formation by lung fibroblasts. (1987). Journal of Biological Chemistry.

31. Omari, S., Makareeva, E., Roberts-Pilgrim, A., Mirigian, L., Jarnik, M., Ott, C., Lippincott-Schwartz, J., and Leikin, S. (2018). Noncanonical autophagy at ER exit sites regulates procollagen turnover. Proc Natl Acad Sci USA 115, E10099–E10108. 10.1073/pnas.1814552115.

32. Ishida, Y., Yamamoto, A., Kitamura, A., Lamandé, S.R., Yoshimori, T., Bateman, J.F., Kubota, H., and Nagata, K. (2009). Autophagic elimination of misfolded procollagen aggregates in the endoplasmic reticulum as a means of cell protection. Mol. Biol. Cell 20, 2744–2754. 10.1091/mbc.E08-11-1092.

33. Forrester, A., De Leonibus, C., Grumati, P., Fasana, E., Piemontese, M., Staiano, L., Fregno, I., Raimondi, A., Marazza, A., Bruno, G., et al. (2019). A selective ER-phagy exerts procollagen quality control via a Calnexin-FAM134B complex. EMBO J. 38. 10.15252/embj.201899847.

34. Reggio, A., Buonomo, V., Berkane, R., Bhaskara, R.M., Tellechea, M., Peluso, I., Polishchuk, E., Di Lorenzo, G., Cirillo, C., Esposito, M., et al. (2021). Role of FAM134 paralogues in endoplasmic reticulum remodeling, ER-phagy, and Collagen quality control. EMBO Rep. 22, e52289. 10.15252/embr.202052289.

35. Bond, A.G., Craigon, C., Chan, K.-H., Testa, A., Karapetsas, A., Fasimoye, R., Macartney, T., Blow, J.J., Alessi, D.R., and Ciulli, A. (2021). Development of BromoTag: A “Bump-and-Hole”-PROTAC System to Induce Potent, Rapid, and Selective Degradation of Tagged Target Proteins. J. Med. Chem. 64, 15477–15502. 10.1021/acs.jmedchem.1c01532.

36. Graham, S.C., Wartosch, L., Gray, S.R., Scourfield, E.J., Deane, J.E., Luzio, J.P., and Owen, D.J. (2013). Structural basis of Vps33A recruitment to the human HOPS complex by Vps16. Proc Natl Acad Sci USA 110, 13345–13350. 10.1073/pnas.1307074110.

37. Yamaguchi, M., Noda, N.N., Yamamoto, H., Shima, T., Kumeta, H., Kobashigawa, Y., Akada, R., Ohsumi, Y., and Inagaki, F. (2012). Structural insights into Atg10-mediated formation of the autophagy-essential Atg12-Atg5 conjugate. Structure 20, 1244–1254. 10.1016/j.str.2012.04.018.

38. Yuan, L., Kenny, S.J., Hemmati, J., Xu, K., and Schekman, R. (2018). TANGO1 and SEC12 are copackaged with procollagen I to facilitate the generation of large COPII carriers. Proc Natl Acad Sci USA 115, E12255–E12264. 10.1073/pnas.1814810115.

39. Liao, Y.-C., Pang, S., Li, W.-P., Shtengel, G., Choi, H., Schaefer, K., Xu, C.S., and Lippincott-Schwartz, J. (2024). COPII with ALG2 and ESCRTs control lysosome-dependent microautophagy of ER exit sites. Dev. Cell 59, 1410–1424.e4. 10.1016/j.devcel.2024.03.027.

40. Chen, S.-Y., Lin, J.-S., and Yang, B.-C. (2014). Modulation of tumor cell stiffness and migration by type IV collagen through direct activation of integrin signaling pathway. Arch. Biochem. Biophys. 555–556, 1–8. 10.1016/j.abb.2014.05.004.

41. Öhlund, D., Franklin, O., Lundberg, E., Lundin, C., and Sund, M. (2013). Type IV collagen stimulates pancreatic cancer cell proliferation, migration, and inhibits apoptosis through an autocrine loop. BMC Cancer 13, 154. 10.1186/1471-2407-13-154.

42. Cui, X., Shan, T., and Qiao, L. (2022). Collagen type IV alpha 1 (COL4A1) silence hampers the invasion, migration and epithelial-mesenchymal transition (EMT) of gastric cancer cells through blocking Hedgehog signaling pathway. Bioengineered 13, 8972–8981. 10.1080/21655979.2022.2053799.

43. Liang, J.R., and Corn, J.E. (2022). A CRISPR view on autophagy. Trends Cell Biol. 32, 1008–1022. 10.1016/j.tcb.2022.04.006.

44. Iavarone, F., Zaninello, M., Perrone, M., Monaco, M., Barth, E., Gaedke, F., Pizzo, M.T., Di Lorenzo, G., Desiderio, V., Sommella, E., et al. (2024). Fam134c and Fam134b shape axonal endoplasmic reticulum architecture in vivo. EMBO Rep. 10.1038/s44319-024-00213-7.

45. Buonomo, V., Lohachova, K., Reggio, A., Cano-Franco, S., Cillo, M., Santorelli, L., Venditti, R., Polishchuk, E., Peluso, I., Brunello, L., et al. (2025). Two FAM134B isoforms differentially regulate ER dynamics during myogenesis. EMBO J. 44, 1039–1073. 10.1038/s44318-024-00356-2.

46. Di Lorenzo, G., Iavarone, F., Maddaluno, M., Plata-Gómez, A.B., Aureli, S., Quezada Meza, C.P., Cinque, L., Palma, A., Reggio, A., Cirillo, C., et al. (2022). Phosphorylation of FAM134C by CK2 controls starvation-induced ER-phagy. Sci. Adv. 8, eabo1215. 10.1126/sciadv.abo1215.

47. Berkane, R., Ho-Xuan, H., Glogger, M., Sanz-Martinez, P., Brunello, L., Glaesner, T., Kuncha, S.K., Holzhüter, K., Cano-Franco, S., Buonomo, V., et al. (2023). The function of ER-phagy receptors is regulated through phosphorylation-dependent ubiquitination pathways. Nat. Commun. 14, 8364. 10.1038/s41467-023-44101-5.

48. González, A., Covarrubias-Pinto, A., Bhaskara, R.M., Glogger, M., Kuncha, S.K., Xavier, A., Seemann, E., Misra, M., Hoffmann, M.E., Bräuning, B., et al. (2023). Ubiquitination regulates ER-phagy and remodelling of endoplasmic reticulum. Nature 618, 394–401. 10.1038/s41586-023-06089-2.

49. Berquez, M., Li, A.L., Luy, M.A., Venida, A.C., O’Loughlin, T., Rademaker, G., Barpanda, A., Hu, J., Yano, J., Wiita, A., et al. (2024). A multi-subunit autophagic capture complex facilitates degradation of ER stalled MHC-I in pancreatic cancer. BioRxiv. 10.1101/2024.10.27.620516.

50. Hashimoto, K., Ochi, H., Sunamura, S., Kosaka, N., Mabuchi, Y., Fukuda, T., Yao, K., Kanda, H., Ae, K., Okawa, A., et al. (2018). Cancer-secreted hsa-miR-940 induces an osteoblastic phenotype in the bone metastatic microenvironment via targeting ARHGAP1 and FAM134A. Proc Natl Acad Sci USA 115, 2204–2209. 10.1073/pnas.1717363115.

51. Gilbert, L.A., Horlbeck, M.A., Adamson, B., Villalta, J.E., Chen, Y., Whitehead, E.H., Guimaraes, C., Panning, B., Ploegh, H.L., Bassik, M.C., et al. (2014). Genome-scale CRISPR-mediated control of gene repression and activation. Cell 159, 647–661. 10.1016/j.cell.2014.09.029.

52. Deutsch, E.W., Bandeira, N., Perez-Riverol, Y., Sharma, V., Carver, J.J., Mendoza, L., Kundu, D.J., Bandla, C., Kamatchinathan, S., Hewapathirana, S., et al. (2025). The ProteomeXchange consortium in 2026: making proteomics data FAIR. Nucleic Acids Res. 10.1093/nar/gkaf1146.

53. Perez-Riverol, Y., Xu, Q.-W., Wang, R., Uszkoreit, J., Griss, J., Sanchez, A., Reisinger, F., Csordas, A., Ternent, T., Del-Toro, N., et al. (2016). PRIDE inspector toolsuite: moving toward a universal visualization tool for proteomics data standard formats and quality assessment of proteomexchange datasets. Mol. Cell. Proteomics 15, 305–317. 10.1074/mcp.O115.050229.

